# Priority conservation areas and a global population estimate for the Critically Endangered Philippine Eagle derived from modelled range metrics using remote sensing habitat characteristics

**DOI:** 10.1101/2021.11.29.470363

**Authors:** Luke J. Sutton, Jayson C. Ibañez, Dennis I. Salvador, Rowell L. Taraya, Guiller S. Opiso, Tristan Luap P. Senarillos, Christopher J.W. McClure

## Abstract

Many range-restricted taxa are currently experiencing population declines yet lack fundamental information regarding distribution and population size. Establishing baseline estimates for both these key biological parameters is however critical for directing conservation planning for at-risk range-restricted species. The International Union for the Conservation of Nature (IUCN) Red List uses three range metrics that define species distributions and inform extinction risk assessments: extent of occurrence (EOO), area of occupancy (AOO) and area of habitat (AOH). However, calculating all three metrics using standard IUCN approaches relies on a geographically representative sample of locations, which for rare species is often spatially biased. Here, we apply model-based interpolation using Species Distribution Models (SDMs), correlating occurrences with remote-sensing covariates, to calculate IUCN range metrics, protected area coverage and a global population estimate for the Critically Endangered Philippine Eagle (*Pithecophaga jefferyi*). Our final range wide continuous SDM had high predictive accuracy (Continuous Boyce Index = 0.927) and when converted to a binary model estimated an AOH = 23,185 km^2^, a maximum EOO = 605,759 km^2^, a minimum EOO = 272,272 km^2^, with an AOO = 53,867 km^2^. Based on inferred habitat from the AOH metric, we estimate a global population of 318 breeding pairs (range: 258-362 pairs), or 636 mature individuals, across the Philippine Eagle global range. Protected areas covered 34 % of AOH, 15 % less than the target representation, with the continuous model identifying key habitat as priority conservation areas. We demonstrate that even when occurrences are geographically biased, robust habitat models can be built that enable quantification of baseline IUCN range metrics, protected area coverage, and a population size estimate. In the absence of adequate location data for many rare and threatened taxa, our method is a promising spatial modelling tool with widespread applications, in particular for island endemics facing high extinction risk.

## Introduction

Species that are rare due to either a restricted geographic range, habitat specificity, or small population size are at greater risk of extinction because their populations may not be as resilient to perturbations in the environment (Rabinowitz *et al*. 1986; Gaston 1994). Therefore, quantifying the two key biological parameters of range extent and population size is fundamental for directing conservation action for threatened rare taxa (Marcer *et al*. 2013; Syfert *et al*. 2014). Habitat loss and fragmentation are the two primary threats to biodiversity globally (Díaz *et al*. 2019), in particular for tropical biodiversity hotspots (Brooks *et al*. 2002). Determining baseline range metrics and population estimates for threatened species with restricted ranges and low abundance can thus inform conservation priorities by quantifying the effects of habitat loss and fragmentation on range extent and population size for these at-risk taxa (IUCN 2001).

The International Union for the Conservation of Nature (IUCN) Red List uses three spatial range metrics that seek to define species distributions and inform extinction risk assessments (IUCN 2019): extent of occurrence (EOO) and area of occupancy (AOO). EOO represents the upper bound of a species distribution, measuring the overall geographic extent of localities and degree of risk spread. Conversely, AOO represents the lower bound of a species distribution. By quantifying where the species actually occurs, AOO is thus linked to population size (Gaston & Fuller 2009). Recently, the IUCN developed a new deductive range metric, area of habitat (AOH, Brooks *et al*. 2019), defined as the extent of habitat factors, such as landcover and elevation, for a species within its range. Estimating AOH is important because it can be used in supporting conservation risk assessments by quantifying habitat loss and protected area coverage (Brooks *et al*. 2019; Sutton *et al*. 2021a).

Protected areas are a fundamental tool for conservation (Rodrigues & Cazalis 2020) and have been successful in reducing habitat loss and fragmentation for many taxa (Brooks *et al*. 2009; Geldmann *et al*. 2013). However, despite wide coverage in the global protected area network, gaps in protected area coverage still exist with new areas being continually added (Rodrigues *et al*. 2004a; 2004b). Additionally, not all protected areas are located in areas deemed effective for conservation, but often designated by socio-economic factors related to competing human activities (Pringle 2017; Morán-Ordóñez 2020). Key Biodiversity Areas (KBAs; BirdLife International 2020), are key sites of international significance for biodiversity which contain: (1) populations of globally threatened species, (2) populations and communities of range or biome restricted species, or (3) substantial congregations of bird species. KBAs also protect areas important for biodiversity and aim to overlap with the entire global protected area network (Donald *et al*. 2019). Identifying key sites within the existing KBA network as new protected areas is usually accomplished using a gap analysis, simultaneously calculating protected area coverage within predicted AOH, thus defining priority sites for protection or conservation action (Scott *et al*. 1993).

Various spatial workflows have been proposed and implemented for calculating AOH, which overlay and clip elevational and landcover preferences within the range of species presence points (Brooks *et al*. 2019). Deductive methods using clipped environmental layers with expert-drawn maps (Harris & Pimm 2008), or inductive modelling methods using inverse distance weighted interpolation (Palacio *et al*. 2021), and logistic regression (Dahal *et al*. 2021; Lumbierres *et al*. 2021), have been successful in estimating AOH but rely on a spatially homogenous sample of presence points. For many rare species in remote, hard to survey areas presence data is either insufficient, or may be heavily biased towards a well-sampled region but lacking elsewhere (Syfert *et al*. 2014; Dahal *et al*. 2021). Because of their rarity, occurrence data for these species is limited and thus calculating range metrics based solely on point data is likely to result in unreliable range metrics (Pena *et al*. 2014). To overcome this issue of sampling bias in calculating AOH a new approach for measuring AOH is required for those rare species with high extinction risk that inhabit remote regions lacking adequate presence data.

The Philippine Eagle (*Pithecophaga jefferyi*) is a large tropical forest raptor and one of the most threatened raptors globally (Bildstein *et al*. 1998), currently classified as ‘Critically Endangered’ on the IUCN Red List (BirdLife International 2018). The Philippine Eagle is endemic to four islands in the Philippine archipelago (Mindanao, Leyte, Samar, and Luzon), sparsely distributed across lowland and montane dipterocarp forests (Salvador & Ibañez 2006). The population has declined drastically over the past 50 years, mainly due to habitat loss through deforestation (Kennedy 1977; Bueser *et al*. 2003; Panopio *et al*. 2021) and persecution (Salvador & Ibañez 2006; Ibañez *et al*. 2016). Thus, the Philippine Eagle fulfils all three components of rarity, and along with its large body size and forest dependency would be associated with a higher risk of extinction (Kittelberger *et al*. 2021). Despite this elevated extinction risk, fundamental aspects of the species biology such as distribution and population size are still uncertain (Collar 1997; Collar *et al*. 1999; BirdLife International 2018) and need updating using a robust methodology.

Most Philippine Eagle research has been conducted on the island of Mindanao (Miranda *et al*. 2000; Bueser *et al*. 2003), and thus occurrence data are biased towards this island. Bueser *et al*. (2003), estimated between 82-233 breeding pairs for Mindanao, and extrapolating this figure across all range islands suggests a global total of between 340 (BirdLife International 2018) and 500 pairs (Salvador & Ibañez 2006). However, pair densities on the other range islands, especially Luzon, are unknown and thus this population size figure should be treated with caution (Miranda *et al*. 2008). Because of these research disparities, there are no current range-wide estimates for the species’ global range extent and population size, despite it being a raptor of high priority for research and conservation (Buechley *et al*. 2019). Indeed, the IUCN Red List suggests that further research into distribution, population size, and ecological requirements is urgently required to inform conservation actions (BirdLife International 2018).

Here, we use Species Distribution Models (SDMs) calibrated with remote sensing covariates and presence-background data for the Philippine Eagle on the island of Mindanao, and then predict into the other less-well sampled islands using inductive model-based interpolation (Rodríguez *et al*. 2007; Franklin 2009). SDMs are predictive spatial models that infer species-habitat associations by correlating species presence points with habitat covariates that represent the focal species optimal conditions and resources (Guisan *et al*. 2017; Matthiopoulos *et al*. 2020).

Indeed, SDMs are able to inform IUCN species range metrics and predict habitat in areas that may lack occurrence data for inclusion in Red List assessments (Marcer *et al*. 2013; Pena *et al*. 2014; Syfert *et al*. 2014; Breiner *et al*. 2017). Using interpolated model predictions, range metrics such as AOH, EOO and AOO can then be calculated based on inferred or predicted habitat following IUCN Red List guidelines (IUCN 2019). First, we present an updated approach to estimating species range metrics and population size based on predicted habitat for the Philippine Eagle, and second, we demonstrate how our methodology can be incorporated into protected area conservation planning for rare species facing extinction.

## Materials and Methods

### Species locations

We compiled Philippine Eagle point localities from the Global Raptor Impact Network (GRIN, McClure *et al*. 2021), a data information system for population monitoring of all raptor species. For the Philippine Eagle, GRIN includes presence-only data consisting of nest locations (*n* = 48) from unstructured surveys (i.e., with no true absence data) conducted on Mindanao by the Philippine Eagle Foundation since 1978 to the present (Miranda *et al*. 2000; Ibañez *et al*. 2016), along with community science data from eBird (*n* = 76 ; Sullivan *et al*. 2009) and the Global Biodiversity Information Facility (*n =* 27; GBIF 2021) (Fig. S1). In addition, we included GPS tracking data from six breeding adult Philippine Eagles from the island of Mindanao (Fig. S2) and pooled this with the nest and community science data to better represent habitat use of a rare species with limited occurrences (Fletcher *et al*. 2019; See Supplementary Material). Duplicate locations and those with no geo-referenced coordinates were removed and then combined into a single range-wide database. A total of 151 geo-referenced records were compiled across the Philippine Eagle range after data cleaning. Only locations recorded from year 1980 onwards were included to match the temporal timeframe of the habitat covariates, whilst retaining sufficient sample size for robust modelling (van Proosdij *et al*. 2016).

For the Mindanao model we used the subset of nest and community science localities from the island of Mindanao, combined with the filtered GPS tracking fixes. We then manually applied a spatial filter between each point, resulting in a single occurrence in each 1-km raster grid cell, resulting in a filtered subset of 435 occurrence records for the Mindanao calibration models. We used spatial filtering because it is the most effective method to account for sampling bias (Kramer-Schadt *et al*. 2013; Boria *et al*. 2014; Fourcade *et al*. 2014) and to ensure we retained the nest locations and GPS fixes as priority data points because of their geolocation accuracy and direct relevance to optimal conditions and resources for Philippine Eagle occurrence. To evaluate the final continuous range-wide model we used all nest and community science localities recorded from 1980 onwards and applied a 1- km spatial filter between each location, regardless of the origin of the point locality.

Applying the 1-km spatial filter resulted in 101 Philippine eagle locations across the entire range for testing calibration accuracy for the final range-wide continuous model.

### Habitat covariates

We defined the species’ accessible area (Barve *et al*. 2011) as consisting of the mainland area of all known range islands: Mindanao, Leyte, Samar, and Luzon (BirdLife International 2018). We extracted the polygons from the World Wildlife Fund (WWF) terrestrial ecoregions shapefile (Olson *et al*. 2001), which correspond to either lowland or montane moist tropical forest. For Luzon, we masked out the tropical pine forest ecoregion in the north of the island because Philippine eagles are habitat specialists of tropical moist dipterocarp forests (Kennedy 1977; Bueser *et al*. 2003; Salvador & Ibañez 2006), and thus unlikely to occur in this ecoregion. Raster covariate layers were cropped to a delimited polygon consisting of the mainland area of all the known range islands. Covariates were selected *a prioiri* based both on environmental factors related empirically to resources and conditions influencing Philippine Eagle distribution (Bueser *et al*. 2003; Ibañez *et al*. 2003; Salvador & Ibañez 2006).

We predicted occurrence using six continuous covariates at a spatial resolution of 30 arc-seconds (∼1-km; Fig. S3) derived from multiple satellite remote sensing products. These consisted of three surface reflectance bands sourced from the Moderate Resolution Imaging Spectroradiometer (MODIS, https://modis.gsfc.nasa.gov/): Band 1 Red (i.e., plant biomass); Band 2 Near Infrared (i.e., leaf and canopy structure); B7 Short Wave Infrared (i.e., senescent biomass), combined with Evergreen Forest landcover downloaded from the EarthEnv repository (https://www.earthenv.org) and a Leaf Area Index biophysical measure downloaded from the Dynamic Habitat Indices repository (https://silvis.forest.wisc.edu/data/dhis/). In addition, we included Human Footprint Index as a measure of human land use sourced from the Socioeconomic Data and Applications Center (SEDAC; https://sedac.ciesin.columbia.edu). Full details on covariates and processing are provided in the Supplementary Material.

### Species Distribution Models

Most Philippine Eagle occurrences and nest locations deposited in GRIN are from the island of Mindanao, with sparse occurrences across the eastern Visayas and Luzon (Fig. S1). Due to this geographical sampling bias, which would likely bias any model predictions (Syfert *et al*. 2014), we developed a model workflow (Fig. S4) to first predict habitat suitability for Mindanao (Fig. S4, box 3a). Next, we then projected each Mindanao model into the islands of the Eastern Visayas and Luzon (Fig. S4, boxes 3b,c), before finally merging each island model into a single range-wide prediction (Fig. S4, box 3c). We parametrized the SDMs using a fine pixel grid (∼1- km), equivalent to fitting an inhomogeneous Poisson process (IPP) with loglinear intensity (Baddeley *et al*. 2010). We did this because the IPP framework is the most effective method to model presence-only data (Warton & Shepherd 2010), common to many raptor monitoring programmes which solely seek to identify occupied areas (Geary *et al*. 2018).

We fitted SDMs using penalized logistic regression, via maximum penalized likelihood estimation (Hefley & Hooten 2015) in the R package maxnet (Phillips *et al*. 2017). Penalized logistic regression imposes a regularization penalty on the model coefficients, shrinking towards zero the coefficients of covariates that contribute the least to the model, reducing model complexity (Gastón & García-Viñas 2011). We limited model complexity because this is necessary when the primary goal is to use SDMs for predictive transferability in space (Helmstetter *et al*. 2020).The maxnet package fits the SDM as a form of infinitely weighted logistic regression (presence weights = 1, background weights = 100), based on the maximum entropy algorithm, MAXENT (Phillips *et al*. 2017). MAXENT is designed for presence-background SDMs and is mathematically equivalent to estimating the parameters for an IPP (Renner & Warton 2013; Renner *et al*. 2015). We used a tuned penalized logistic regression algorithm because this approach outperforms other SDM algorithms (Valavi *et al*. 2021), including ensemble averaged methods (Hao *et al*. 2020). Full details on the model parameter settings are outlined in the Supplementary Material.

We evaluated calibration accuracy for the Mindanao model using a random sample of 4,350 background points at a recommended 1:10 ratio to the presence data (Helmstetter *et al*. 2020). For the range-wide model we used a random sample of 10,000 background points as pseudo-absences recommended to sufficiently sample the background calibration environment (Barbet-Massin *et al*. 2012; Guevara *et al*. 2018). We used Continuous Boyce index (CBI; Hirzel *et al*. 2006) as a threshold- independent metric of how predictions differ from a random distribution of observed presences (Boyce *et al*. 2002). CBI is consistent with a Spearman correlation (*r_s_*) and ranges from -1 to +1. Positive values indicate predictions consistent with observed presences, values close to zero suggest no difference with a random model, and negative values indicate areas with frequent presences having low environmental suitability. Mean CBI was calculated using five-fold cross-validation on 20 % test data with a moving window for threshold-independence and 101 defined bins in the R package enmSdm (Smith 2019).

For the Mindanao model, we further tested the optimal predictions against random expectations using partial Receiver Operating Characteristic ratios (pROC), which estimate model performance by giving precedence to omission errors over commission errors (Peterson *et al*. 2008). Partial ROC ratios range from 0 to 2 with 1 indicating a random model. Function parameters were set with a 10% omission error rate, and 1000 bootstrap replicates on 50% test data to determine significant (*α* = 0.05) pROC values >1.0 in the R package ENMGadgets (Barve & Barve, 2013).

Lastly, the final range-wide continuous prediction was tested using CBI and then converted into a binary threshold prediction based on expert validation from J.C.I., which we term *model* AOH (Fig. S4, box 5), so as to be distinct from the standard IUCN AOH methodology (Brooks *et al*. 2019).

We validated our models in conjunction with expert judgement because this approach gives most benefit to conservation risk assessments (Marcer *et al*. 2013; Syfert *et al*. 2014). Following modelling protocols established by Velásquez-Tibatá *et al*. (2019), we assessed a range of four binary thresholds for biological realism (median, 75 % upper quantile, maximizing the sum of sensitivity and specificity (maxTSS) and Cohen’s Kappa), using expert critical feedback to assess the predictive ability of our models (Fig. S4, boxes 4b,c). Both maxTSS and upper quantile binary models were evaluated as plausible range extents but we opted for maxTSS because this threshold is recommended for spatial conservation applications (Liu *et al*. 2013). We followed a participatory modelling process methodology to ensure a robust expert validation of our models, concurring with current knowledge of species biology and its application to conservation planning (Ferraz *et al*. 2020).

### Range sizes

To calculate *model* AOH in suitable pixels we reclassified the continuous prediction to a binary threshold prediction (Fig. S4, boxes 4a,b), using all pixel values equal to or greater than the maxTSS threshold from the continuous model. We calculated two further IUCN range metrics from our *model* AOH binary prediction. First, Area of Occupancy (AOO) was calculated as the number of raster pixels predicted to be occupied, scaled to a 2x2 km grid (4-km^2^ cells) following IUCN guidelines (IUCN 2018) in the R package redlistr (Lee *et al*. 2019). Second, we converted the *model* AOH raster to a polygon using an 8-neighbour patch rule and applied a smoothing function using the Chaikin algorithm (Chaikin 1974) in the R package smoothr (Strimas-Mackey 2021). From this we calculated Extent of Occurrence (EOO), fitting a minimum convex polygon (MCP) around the furthest boundaries of the smoothed *model* AOH polygon following IUCN guidelines (IUCN 2018). We calculated both a maximum EOO, including all the area with the MCP, and a minimum EOO, masking out the areas that could never be occupied within the MCP, in our case over the ocean (Mercer *et al*. 2013). All range metric calculations were performed using a Transverse cylindrical equal area projection following IUCN guidelines (IUCN 2018).

### Population size estimation

We calculated the number of Philippine Eagle pairs our *model* AOH could support as directly proportional to the available habitat within a given home range required by a breeding pair of Philippine Eagles (Kennedy 1977; Krupa 1989). Based on the premise that central-place foragers, such as the Philippine Eagle, require a semi-fixed area of habitat to survive and reproduce, we calculated the habitat area required for each pair on home range estimates from six breeding adult Philippine Eagles fitted with satellite telemetry tags (Table S1). We calculated home range sizes using three different estimators to provide a range of habitat area estimates for calculating population size because of variation in outputs between different home range estimation methods (Signer & Fieberg 2021; See Supplementary Material).

Using the habitat area from the three estimates, we then calculated the median, and a range of minimum to maximum population sizes of potential breeding pairs that our *model* AOH prediction could support using the formulation of Kennedy (1977),

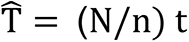

where T*hat* = total population size; N = area of habitat; n = home range estimate and t = sample total multiplied by 2. We used the IUCN Red List definitions for population size as the total number of mature individuals across the species range (IUCN 2019), then divided that figure by 2 to give the number of potential breeding pairs.

### Protected Area Gap Analysis

We assessed the level of protected area coverage within the Key Biodiversity Area (KBA) network using the World Database of Protected Area (WDPA) terrestrial shapefile for the Philippines (as of June 2021; UNEP-WCMC & IUCN 2021). We quantified how much protected area representation is needed for the Philippine Eagle dependent on the *model* AOH to calculate a protected area ‘representation target’ following the formulation of Rodrigues *et al*. (2004a),

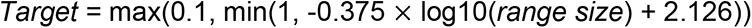

where ‘*Target’* is equal to the percentage of protected target representation required for the species ‘*range size’*, as used in subsequent applications of the formula (Butchart *et al*. 2015; Di Marco *et al*. 2017). We calculated the difference between the current level of KBA coverage compared to the target level representation for terrestrial WDPA coverage using the *model* AOH intersected with the KBA polygons (as of September 2020; BirdLife International 2020), establishing those KBAs covering areas of habitat suitability ≥ maxTSS threshold. The KBA network polygons were then overlaid with the continuous maps for each island identifying gaps in habitat suitability ≥ maxTSS threshold which were not covered by the terrestrial WDPA polygons. We used the continuous models to identify priority conservation areas because continuous predictions give more precision for identifying spatial conservation planning hotspots than binary outcomes (Guillera-Arroita *et al*. 2015). Model development and geospatial analysis were performed in R (v3.5.1; R Core Team, 2018) using the raster (Hijmans 2017), rgdal (Bivand *et al*. 2019), rgeos (Bivand & Rundle 2019) and sp (Bivand *et al*. 2013) packages.

## Results

### Species Distribution Models

For the Mindanao models, six candidate models had an ΔAIC_c_ ≤ 2, and the model with the highest regularization multiplier penalty (β) that used both linear and quadratic terms with the lowest omission rate was selected (Model 4 in Table S2). The best-fit SDM for the island of Mindanao (ΔAIC_c_ = 1.665) had a beta coefficient penalty of β = 1.5 with linear and quadratic terms as model parameters, with high calibration accuracy (mean CBI = 0.947), and was robust against random expectations (pROC = 1.558, SD± 0.068, range: 1.314 – 1.797). The optimal model shrinkage penalty was able to retain ten non-zero beta coefficients, only setting to zero the quadratic terms for Band 1 Red and Evergreen Forest (Table 1), meaning most covariate terms were highly informative to model prediction (Figs. S5-S7).

**Table 1.** Parameter estimates from the penalized beta coefficients derived from the linear and quadratic response functions for each habitat covariate from the optimal Species Distribution Model for the Philippine Eagle on Mindanao island.

| Covariate | Linear | Quadratic |
| --- | --- | --- |
| B1 Red | 5.707 | 0.000 |
| B2 Near Infrared | -6.851 | -6.566 |
| B7 Short Wave Infrared | 4.624 | 8.621 |
| Evergreen Forest | 0.043 | 0.000 |
| Human Footprint Index | 0.062 | -0.002 |
| Leaf Area Index | 1.254 | -0.413 |

From the penalized beta coefficients (Table 1), Philippine Eagles on Mindanao were most positively associated with Band 1 Red surface reflectance values > 0.4 (i.e., dense, healthy green leaf biomass), followed by lower Band 7 Short Wave Infrared surface reflectance values between 0.2-0.4 (i.e., senescent or old-growth biomass) and most negatively associated with Band 2 Near Infrared surface reflectance values > 0.3 (i.e., leaf and canopy structure; Fig. 1). Philippine Eagles had a unimodal response to Leaf Area Index values of 1.5 (i.e., multiple layered canopy cover), and a positive linear response to Evergreen Forest cover between 70-80 % (Fig. 1).

**Figure 1.**
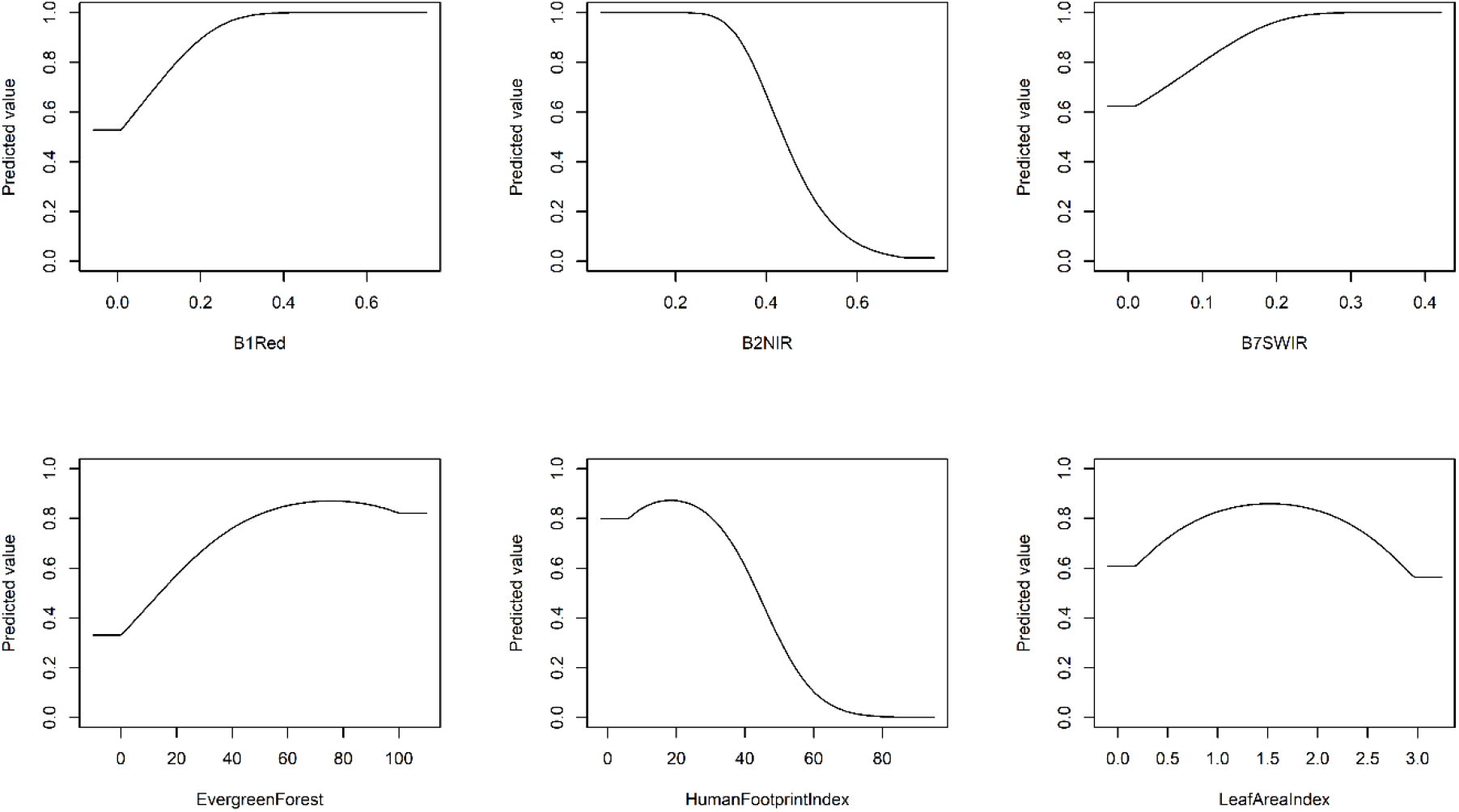
Penalized logistic regression response functions for each habitat covariate from the optimal Species Distribution Model for the Philippine Eagle on Mindanao island. The curves show the contribution to model prediction (y-axis) as a function of each continuous habitat covariate (x-axis). Maximum values in each response curve define the highest predicted relative suitability. The response curves reflect the partial dependence on predicted suitability for each covariate and the dependencies produced by interactions between the selected covariate and all other covariates. Absolute reflectance values on the x-axes of the top row panels are expressed as the ratio of reflected over incoming radiation, meaning reflectance can be measured between the values of zero and one. Reflectance values of 3-4 indicate healthy vegetation.

Philippine Eagles had a positive relationship up to Human Footprint Index values of 20 (i.e., areas of low human impact) decreasing sharply to areas with high impact human infrastructure.

On Mindanao, the largest continuous areas of Philippine Eagle habitat were confined to mountainous areas with high forest cover across the eastern and central mountain ranges of Kitanglad, Pantaron, Diwata, and the Bukidnon plateau (Fig. 2). Patchy areas of habitat were identified throughout western Mindanao, largely confined to areas of steep, forested terrain, and extending further south into the Tiruray Highlands and Mount Latian complex. Little habitat was predicted across the now largely deforested lowland plains. The range-wide continuous model had high predictive performance (CBI = 0.927) and was able to capture key areas of habitat when projected to the islands of Luzon and the Eastern Visayas (Fig. 3). For the Eastern Visayas, highest habitat suitability was predicted in a small area of north- eastern Samar. Only small patches of high suitability habitat were predicted for Leyte. In Luzon the largest continuous area of Philippine Eagle habitat was predicted in the northern Sierra Madres mountain range in the east of the island, with smaller patches further south. Further high suitability habitat was predicted in the north of Luzon in the northern Cordillera mountain range and a smaller area of habitat was predicted for the Zambales mountain range in the far west of Luzon.

**Figure 2.**
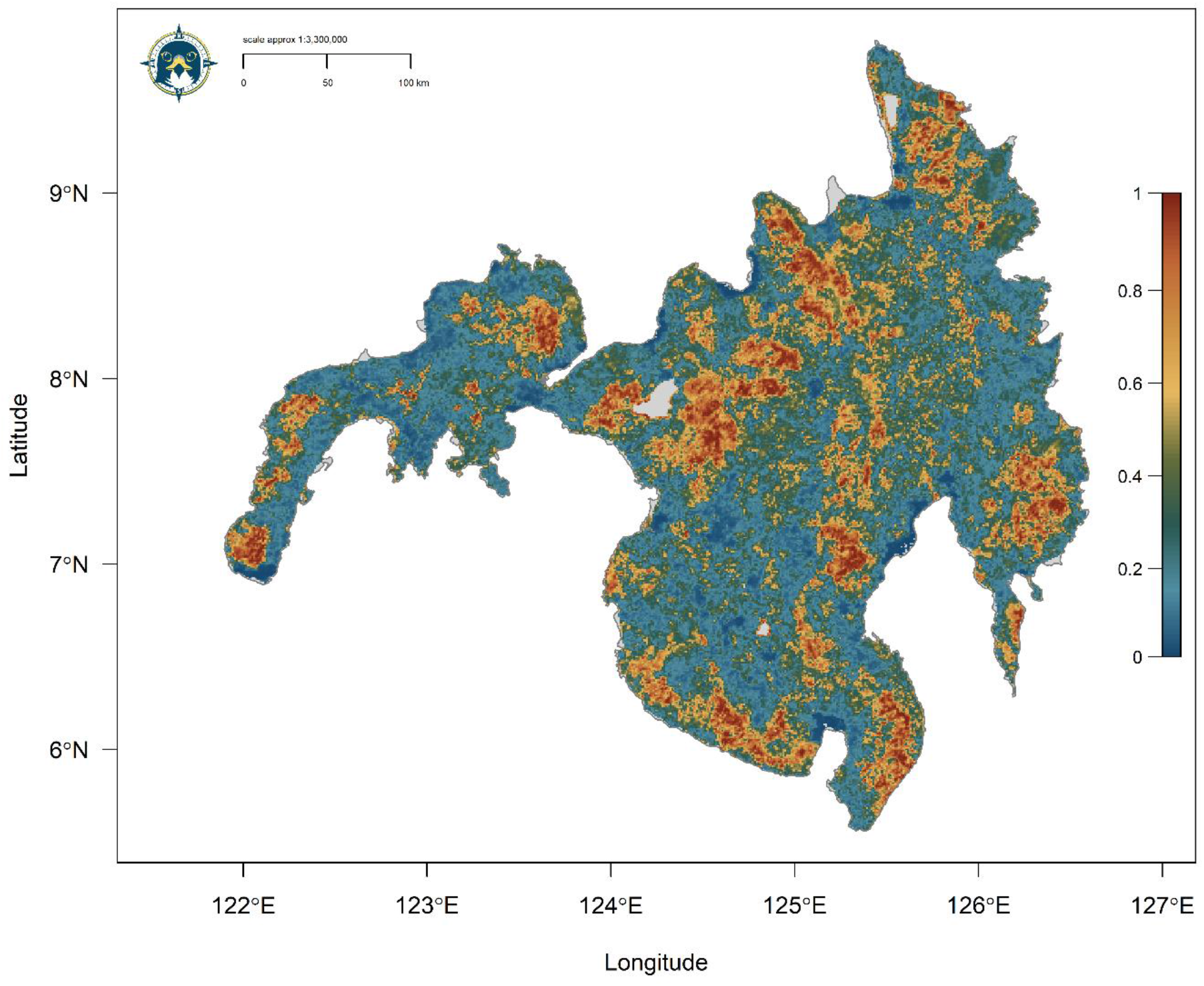
Continuous Species Distribution Model for the Philippine Eagle on the island of Mindanao using a penalized logistic regression model algorithm. Map denotes habitat suitability prediction with red areas (values closer to 1) having highest habitat suitability, orange/yellow moderate suitability and blue/green low suitability (values closer to zero).

**Figure 3.**
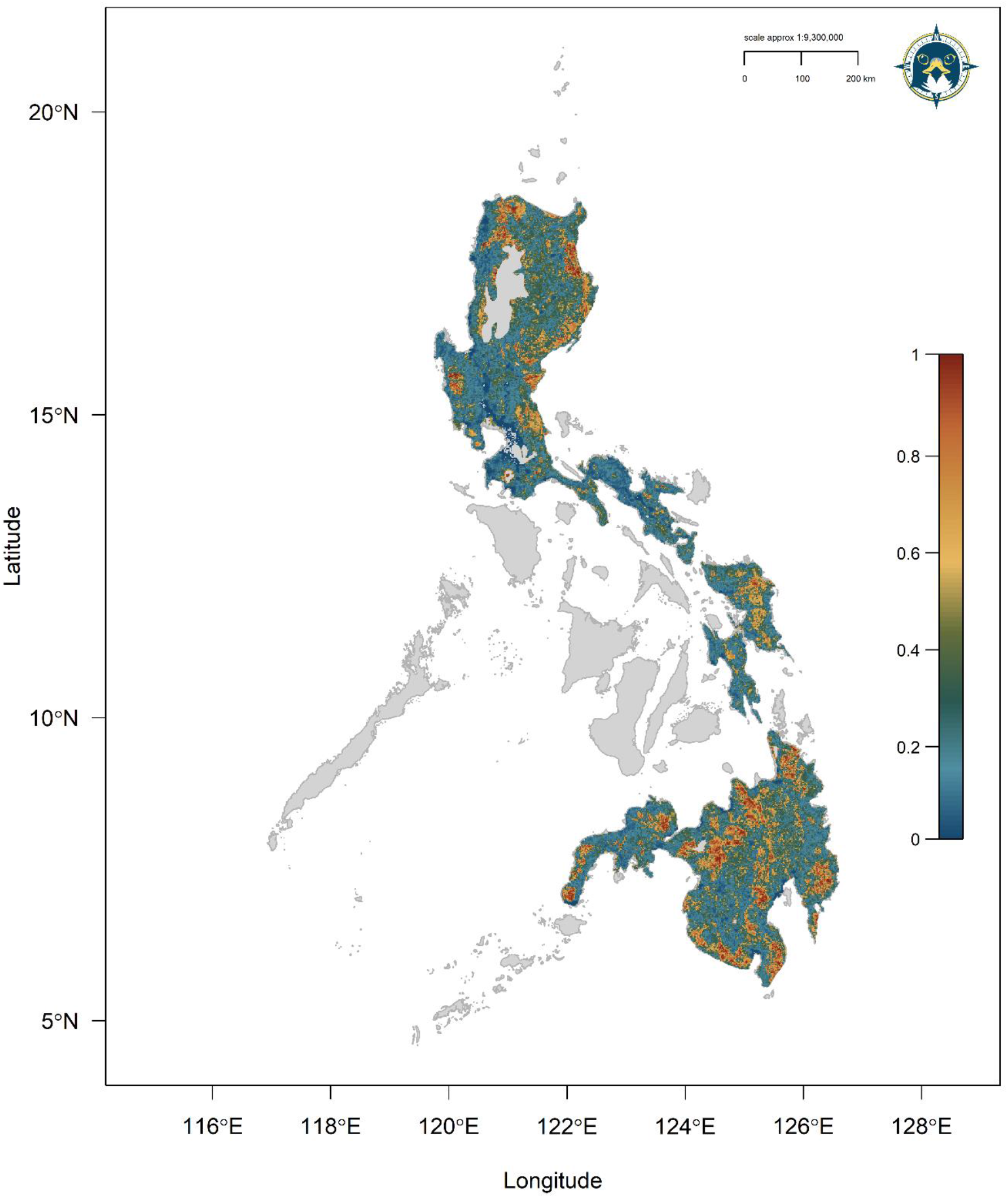
Range-wide Species Distribution Model for the Philippine Eagle using a penalized logistic regression model algorithm. Map denotes continuous prediction with red areas (values closer to 1) having highest habitat suitability, orange/yellow moderate suitability and blue/green low suitability.

### Range metrics and population size

The reclassified binary model (maxTSS threshold = 0.620) calculated a *model* AOH = 23,185 km^2^ (Fig. 4). From the *model* AOH, maximum EOO was 605,759 km^2^ and minimum EOO 272,272 km^2^ (Fig. 4), with an AOO = 53,867 km^2^. **T**he median territorial habitat area based on the home range estimates from the six adults was 73 km^2^ using the KDE estimator (Table 2; Fig. S8), with a minimum and maximum range of 64 and 90 km^2^ of territorial habitat area using the median home range estimates from the r-LoCoH and 99 % MCP estimators respectively (Table 2; Fig. S9). Using our formulation based on habitat area from home range estimates, we calculated the *model* AOH could potentially support 318 breeding pairs (range: 258- 362), or 636 mature individuals, across the entire Philippine Eagle range based on the *model* AOH area of 23,185 km^2^. The area of habitat in Mindanao (14,315 km^2^) could potentially support 196 breeding pairs (range: 159-224; Fig. S10), in Luzon (AOH = 7,632 km^2^) 105 pairs (range: 85-119; Fig. S11) and in the Eastern Visayas (AOH = 1,238 km^2^) 17 pairs (range: 14-20; Fig. S12).

**Figure 4.**
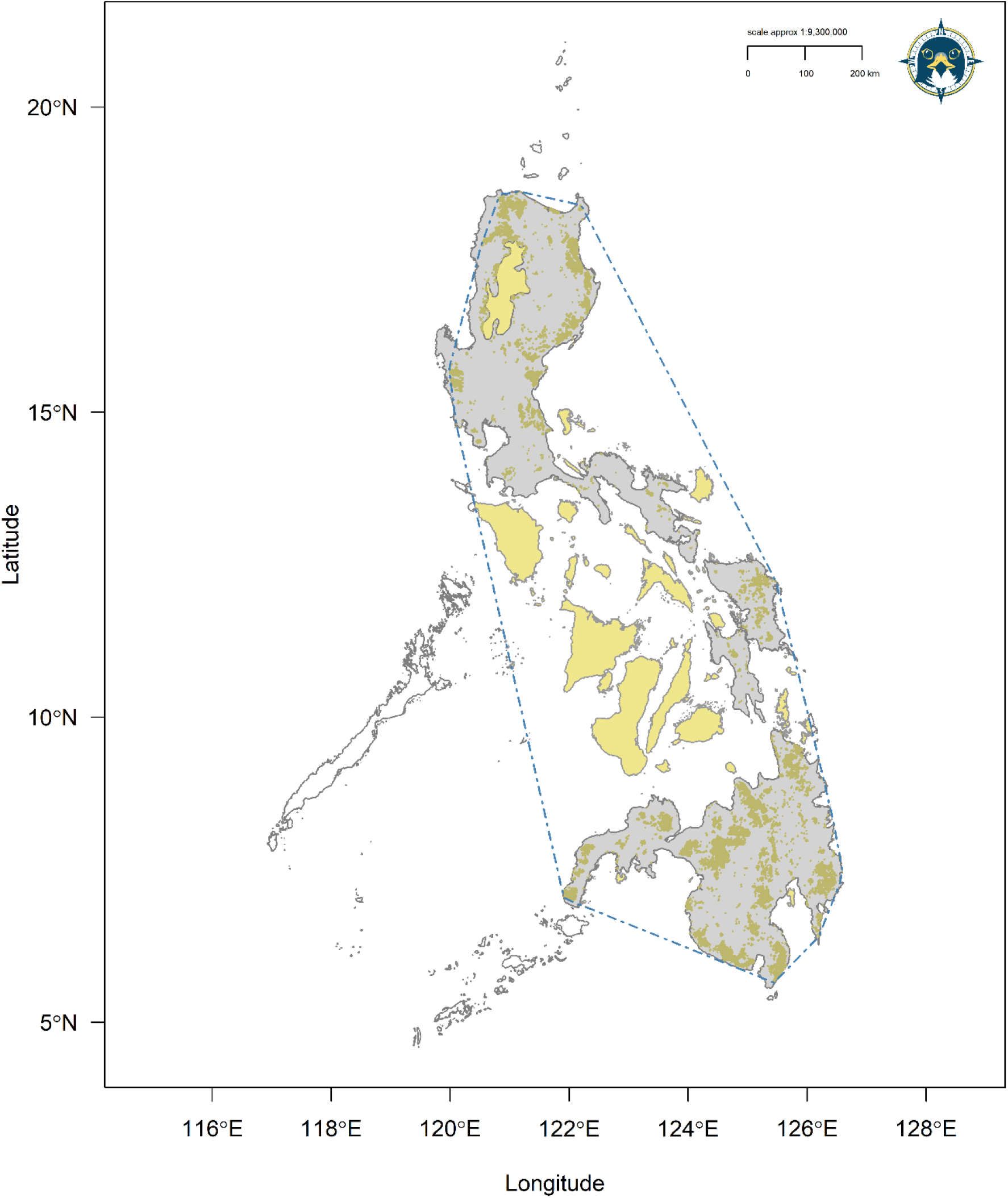
Range metrics for the Philippine Eagle showing the reclassified binary *model* area of habitat (AOH) area (brown) and extent of occurrence (EOO, hashed blue polygon). Grey island polygons represent the species accessible area. Yellow polygons define the national boundary of the Philippines not within the species accessible area.

**Table 2.** Home range estimates for six breeding adult Philippine Eagles using three home range estimators. r-LoCoH = radius Local Convex Hull; KDE = Kernel Density Estimate, MCP = Minimum Convex Polygon. All values are km^2^.

| Adult ID | r-LoCoH | 95% KDE | 99% MCP |
| --- | --- | --- | --- |
| 001F | 61 | 70 | 88 |
| 002F | 85 | 75 | 105 |
| 003F | 37 | 43 | 36 |
| 004M | 66 | 107 | 91 |
| 005M | 53 | 41 | 57 |
| 006F | 120 | 147 | 173 |
| Median | 64 | 73 | 90 |

### Priority conservation areas

Across the Philippine Eagle range, the current WDPA network covered 34.2 % (7,936 km^2^) of the *model* AOH (Fig. S13), 14.8 % less than the target protected area representation of 49 %, with the KBA network covering 71 % (16,471 km^2^) of the *model* AOH (Fig. S14), more than double the coverage within the WDPA network.

Priority areas of Philippine Eagle habitat which are currently classified as KBAs but without protected area coverage in the WDPA network were identified on all range islands.

On Mindanao, priority KBAs for upgrading to protected areas include (Fig. 5): (1) Mount Hilong-hilong and (2) Mount Kampalili-Putting Bato in the Eastern Mindanao Biodiversity Corridor. In southern-central Mindanao, priorities are extending the protected area for Mount Apo Natural Park (3) into the northern part of the KBA, along with protected status for the Mount Latian complex and Mount Busa-Kiamba KBAs (4). Protected areas could also be extended in the Mount Piagayungan and Butig Mountains and Munai/Tambo KBAs (5) in east-central Mindanao. In northern-central Mindanao, priority KBAs for protection include the Mount Kaluayan – Mount Kinabalian Complex along with the adjacent Mount Balatukan, and the Mount Tago Range KBAs (6). In addition we recommend new KBAs and/or protected areas be established in: (A) Sibuco-Sirawai region of western Mindanao (Fig. 5; dashed blue circle A); (B) the Daguma Range-Palimbang region of southern Mindanao (dashed blue circle B), and (C) Mount Sinaka in central Mindanao (dashed blue circle C).

**Figure 5.**
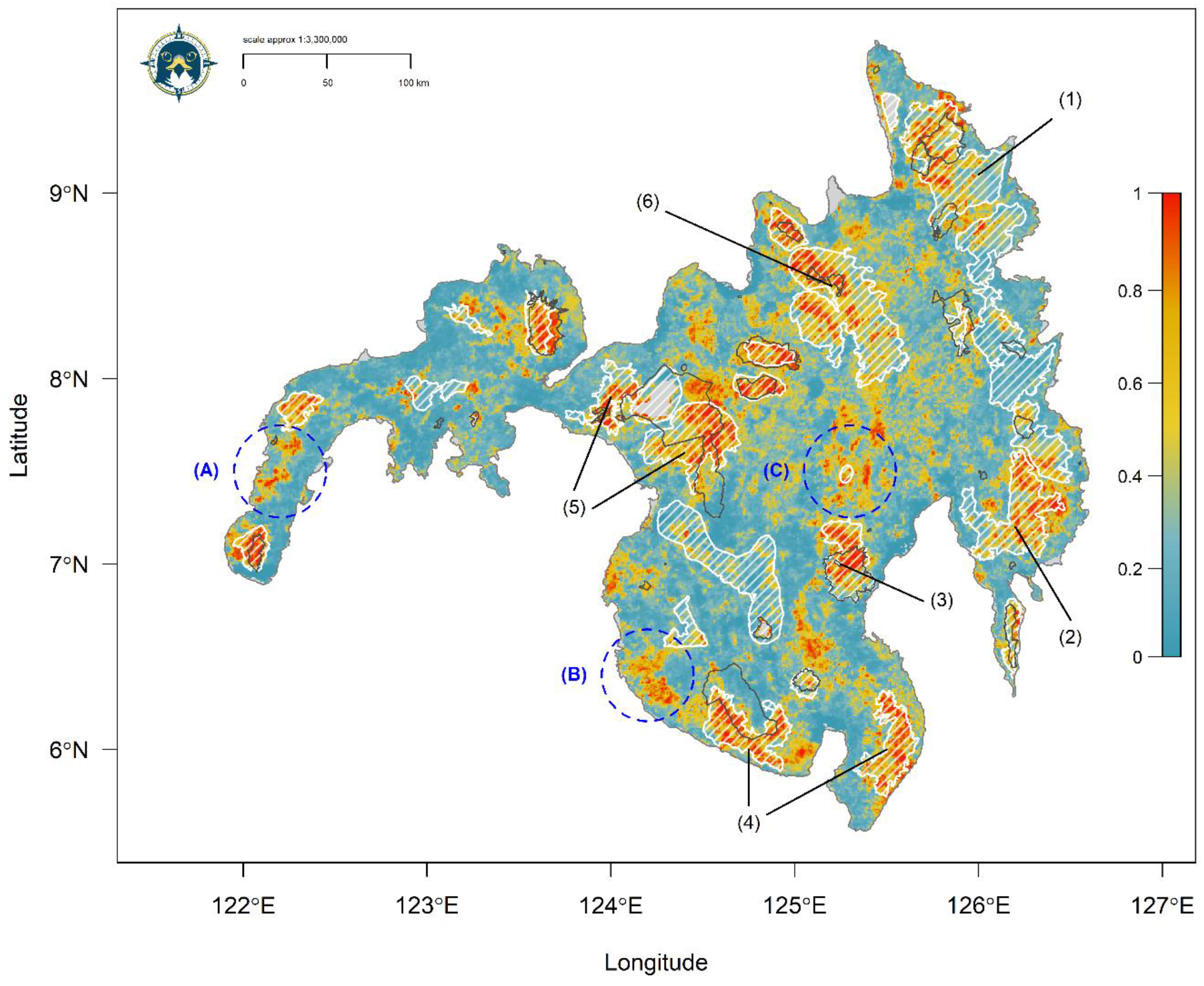
Gap Analysis for Philippine Eagle habitat on the island of Mindanao showing spatial coverage of the World Database on Protected Areas (WDPA) network (dark grey polygons) compared to the Key Biodiversity Area (KBA) network coverage (white hashed polygons) within the continuous model prediction. Map denotes habitat suitability prediction with red areas (values closer to 1) having highest habitat suitability, yellow moderate suitability and blue low suitability (values closer to zero). Numbered arrows indicate priority KBAs for protection: (1) Mount Hilong-hilong, (2) Mount Kampalili-Putting Bato, (3) Mount Apo, (4) Mount Latian & Mount Busa-Kiamba, (5) Mount Piagayungan & Butig Mountains and Munai/Tambo, (6) Mount Kaluayan-Mount Kinabalian Complex, Mount Balatukan, and Mount Tago Range. Hashed blue circles indicate areas og high suitability habitat recommended as new KBAs and/or proected areas: (A) Sibuco-Sirawai, (B) Daguma Range-Palimbang, and (C) Mount Sinaka.

In the Eastern Visayas, most habitat within the KBA network on Samar was contained within the Samar Island Natural Park (IUCN Cat. II; Fig. 6). The north-east of the island had highest habitat suitability and should be prioritized for further protection extending across north-east Samar beyond the national park to include high suitability habitat which has no coverage within either the KBA or protected area networks. The priority KBA for protection in Leyte was Anonang-Lobi Range (Fig. 6). In addition, there were small pockets of high suitability habitat in the east of Leyte outside the current Anonang-Lobi Range KBA which could be extended for further habitat protection and potential reintroductions if given protected status. For Luzon, priority KBAs for proposed new protected areas include (Fig. 7): (1) the Apayao Lowland Forest in northern Luzon, along with extending this KBA and the Balbalasang-Balbalan KBA west to cover further high suitability habitat. Connecting high suitability habitat along the Sierra Madre Range by protecting the North Central Sierra Madre Mountains KBA (2) and Mount Dingalan and Aurora Memorial National Park KBAs (3) in eastern Luzon. Lastly, (4) the Zambales Mountains could also be upgraded for protection if surveys identify a population here, otherwise the KBA should be prioritized for potential reintroductions.

**Figure 6.**
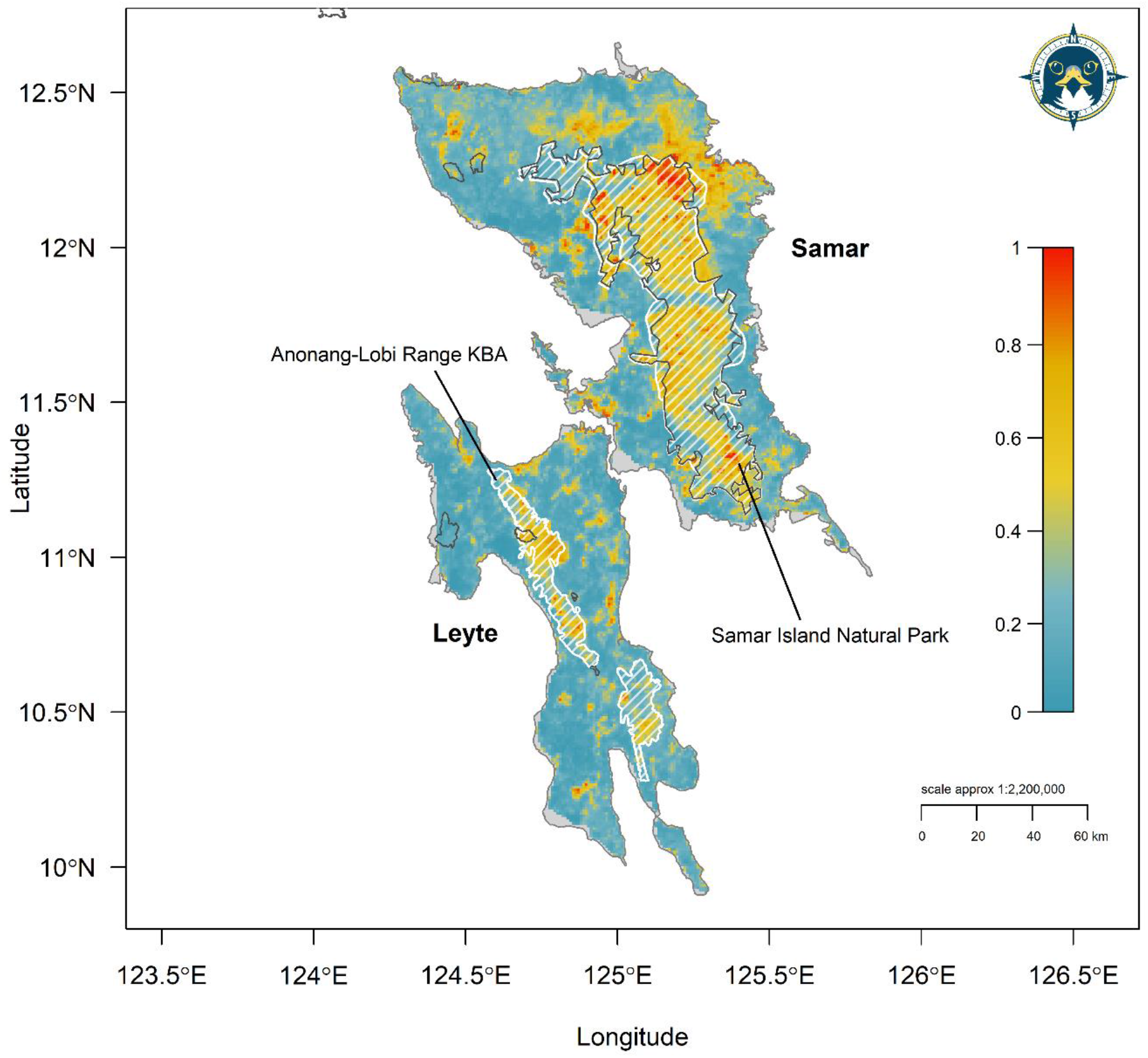
Gap Analysis for Philippine Eagle habitat on the islands of Leyte and Samar in the Eastern Visayas showing spatial coverage of the World Database on Protected Areas (WDPA) network (dark grey polygons) compared to the Key Biodiversity Area (KBA) network coverage (white hashed polygons) within the continuous model prediction. Map denotes habitat suitability prediction with red areas (values closer to 1) having highest habitat suitability, yellow moderate suitability and blue low suitability (values closer to zero).

**Figure 7.**
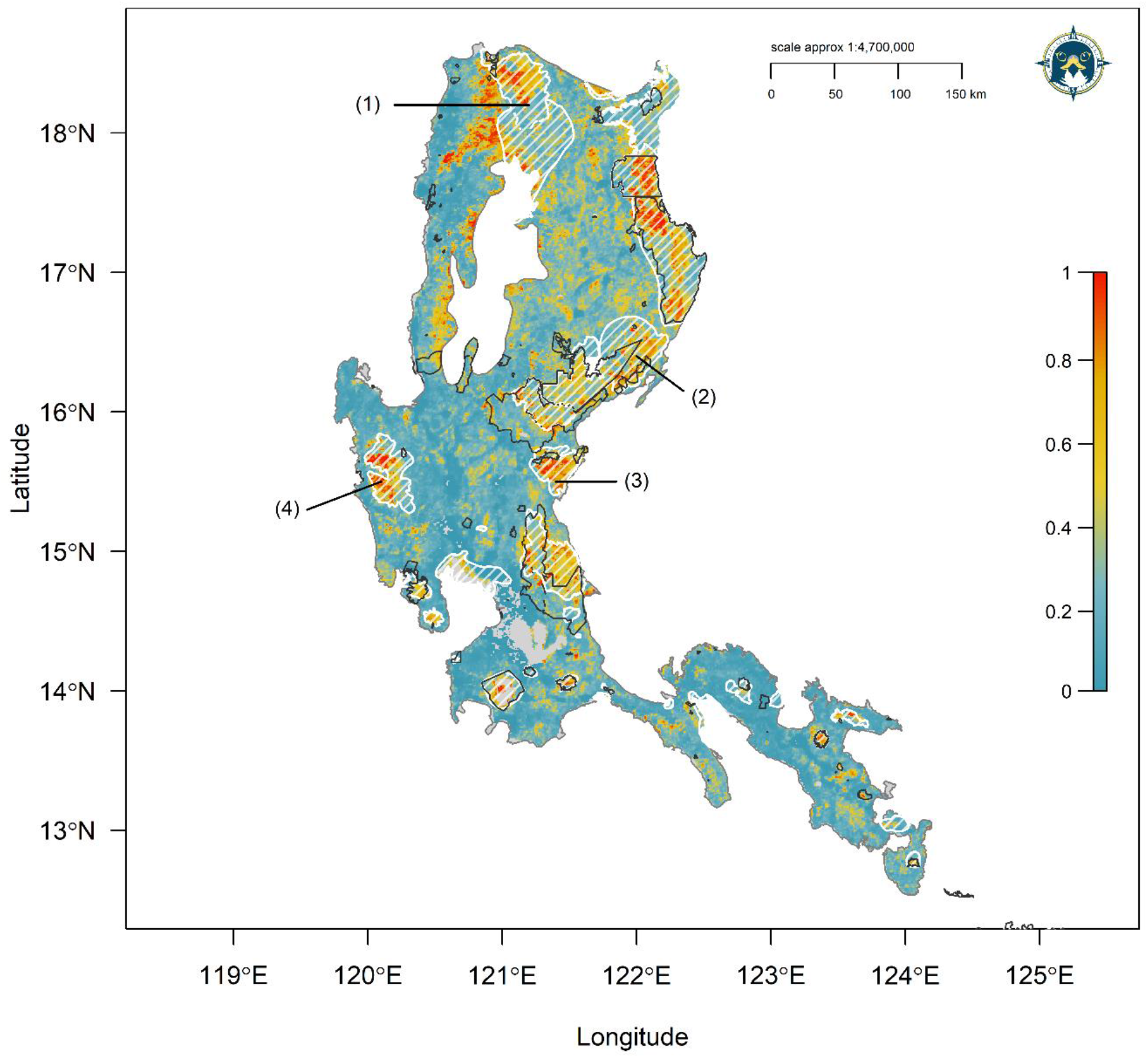
Gap Analysis for Philippine Eagle habitat on the island of Luzon showing spatial coverage of the World Database on Protected Areas (WDPA) network (dark grey polygons) compared to the Key Biodiversity Area (KBA) network coverage (white hashed polygons) within the continuous model prediction. Map denotes habitat suitability prediction with red areas (values closer to 1) having highest habitat suitability, yellow moderate suitability and blue low suitability (values closer to zero). Numbered arrows indicate priority KBAs for protection: (1) Apayao Lowland Forest and Balbalasang- Balbalan, (2) North Central Sierra Madre Mountains, (3) Mount Dingalan and Aurora Memorial National Park, (4) Zambales Mountains.

## Discussion

Range-restricted tropical raptors are particularly threatened by human-induced land use activities (Cruz *et al*. 2021), with many experiencing severe population declines and in need of immediate research and conservation (McClure *et al*. 2018; Buechley *et al*. 2019). Correlating occurrence data from multiple sources with remote-sensing environmental data, we provide a first estimate of Area of Habitat for the Philippine Eagle, update the species’ IUCN range metrics, and provide a baseline global population estimate. By establishing baselines for these key biological parameters, we then applied our model outputs for directing long-term monitoring and priority conservation planning for this Critically Endangered raptor. Despite issues of geographic sampling bias in our occurrence dataset, we were able to overcome any analytical setbacks by implementing a robust and straightforward modelling framework. We view our methodology as a widely applicable tool for quantifying species-habitat associations for many taxa of conservation concern.

Our *model* AOH map updates previous estimates of potential habitat for the Philippine Eagle, further refining the habitat map from Krupa (1989). Our *model* AOH estimate of 23,185 km^2^ confirms the Philippine Eagle as a range-restricted and endemic species, which are not always mutually exclusive (Gaston 1994). We were able to use our binary model prediction to calculate a first estimate for AOO (53,867 km^2^) and an updated EOO bounded from the *model* AOH polygon (see Fig. 4). Our maximum EOO (605,759 km^2^), was 9 % larger than the current IUCN estimate (551,000 km^2^; BirdLife International 2018). However, when considering the area of EOO not covering the unoccupiable area of the ocean, our minimum EOO (272,272 km^2^) was 50 % less. We posit that using a minimum EOO is more relevant for species that range across island archipelagos because including areas that cannot be occupied within the entire area of the MCP in the EOO range metric calculation is potentially misleading. We recognise the need to have a consistent global methodology for species range metrics but not at the cost of inflating risk spread in the EOO range metric for threatened island ranging species. Thus, we recommend that both a minimum and maximum EOO be reported in future IUCN range assessments where relevant.

Area of Habitat maps are useful in many conservation applications such as protected area assessments, targeting surveys and monitoring habitat loss (Brooks *et al*. 2019). Here, we also applied our *model* AOH to calculating a key biological parameter used in IUCN conservation risk assessments, that of a global population estimate (IUCN 2019). However, we stress that our global estimate of 318 pairs (636 mature individuals) is the *potential* breeding population size based on inferred habitat from SDM outputs which may not always link to population parameters (Lee-Yaw *et al*. 2021). Our global population size of 318 pairs was only slightly less than the current IUCN estimate of 340 pairs (BirdLife International 2018) but greater than an earlier estimate of 88-221 pairs (Krupa 1989). However, the key difference here is that we used an empirical estimate of habitat area needed for each pair based on home range estimates. Assuming our baseline population estimate is accurate, we urge more investment and research, such as ground truthing surveys, into conserving these remaining populations and their forest habitat.

Our median population estimate for Mindanao (*n* = 196 pairs) was within the range of the current estimate for the island (82-233 pairs; Bueser *et al*. 2003), but greater than other previous population estimates (Kennedy 1977; Krupa 1989). Bueser *et al*. (2003) calculated population size using a different method based on habitat within a circular plot around known nest sites from nearest neighbour distances and estimated remaining total forest habitat. That our population size estimate for Mindanao was within the range given by Bueser *et al*. (2003), gives credence to our method that uses home range estimates with our area of habitat size. The median population estimate for Luzon (*n* = 105 pairs) was nearly half that for Mindanao but higher than from a previous estimate of 33-83 pairs, that used assumed territory sizes of 60-100 km^2^ with the then area of remaining forest habitat in the Philippines (Krupa 1989). Exploratory ground truthing surveys are required across Luzon to establish the accuracy of our baseline population estimate.

Historically, Philippine Eagles were recorded throughout Luzon (Kennedy 1977) but with most records largely restricted to the Sierra Madre range (Poulsen 1995; Panopio *et al*. 2021), albeit at assumed low densities (Krupa 1989). Indeed, recent surveys in the north of Luzon discovered the first nest in the northern Cordillera range (Abaño *et al*. 2016), with our model predicting extensive Philippine Eagle habitat across both the Sierra Madre and Cordillera ranges. Interestingly, our estimate of 17 pairs (range: 14-20 pairs) for the Eastern Visayan islands of Leyte and Samar was similar to earlier estimates (Kennedy 1977; Krupa 1989). Previous pair numbers for Samar ranged between 8-19 pairs (Krupa 1989), with numbers on Leyte estimated to be between 8-10 pairs (Kennedy 1977), or as low as 1-4 pairs (Krupa 1989). Similar to Luzon, we urge more surveys on Leyte and Samar to ground truth our estimates.

The Philippines is one of the most biodiverse countries globally (Myers *et al*. 2000), with an established community-based protected area system (Senga 2001; Posa *et al*. 2008). Our gap analysis was able to identify 15 current KBAs on all range islands (both with and without any form of protection), as priority sites for new or extended protected areas within the current network, along with two priority sites for reintroductions on Leyte and Luzon and three recommended sites for new protected designations on Mindanao. Due to the Philippine Eagle’s reliance on tropical dipterocarp forest, we recommend designating these KBAs as new protected areas, Indigenous and Community Conserved Areas (ICCAs), and/or Local Conservation Areas(LCAs), thus connecting the remaining habitat patches which is key to the species future survival (Poulsen 1995; Posa *et al*. 2008). Further, protecting these key areas of tropical forest habitat should also be beneficial for prioritizing reintroduction sites. A key advantage of using covariates derived from MODIS satellite remote sensing data means constant monitoring can be established for changes in vegetation (Perez & Comiso 2014). New MODIS covariates can then be used in updated models for the key biological and conservation parameters for area of habitat, population size, and protected area coverage, meaning rapid action can tackle emerging threats when needed.

There is no one overriding ‘best’ method for modelling species-habitat associations but multiple approaches dependent on the purpose of the study (Qiao *et al*. 2015). Our approach was useful because of its ability to predict beyond the known range limits of the Philippine Eagle, providing a *potential* area of habitat (*sensu* Sutton *et al*. 2021a,b). This was appropriate in this context when our goal was to provide baseline estimates for global range extent and population size, with geographically biased species locations, rendering standard habitat modelling approaches unsuitable. Further, the standard IUCN approach to estimating AOH uses solely landcover and elevation as covariates (Brooks *et al*. 2019). Here, along with landcover we also incorporated important predictors for determining species’ habitat associations such as those from raw surface reflectance values and human land use (Guisan *et al*. 2017), which improved model predictions compared to an initial model using climate and landcover (Sutton *et al*. 2021c). We recommend that analysts consider remote sensing variables in future area of habitat assessments to fully capture the environmental range limits for a given taxa.

Whilst we envision broad applications for our methodology, we recognise that our spatial workflow is likely most useful for island endemic species with low numbers of occurrences, or with pronounced geographic sampling bias in species locations.

Despite potential issues with sampling bias from pooling occurrences from disparate data sources (Fletcher *et al*. 2019), we were able to use the spatial filter to account for sampling bias and use pooled data because we had no true absences to use in our models, only presences. We sampled pseudo-absences from our study area but the assumption that all absences would be within that study area (thus approximating the model integral) is difficult to assess (Hefley & Hooten 2016).

Rectifying the issues for this form of sampling bias with an appropriate data model is currently unknown (Hefley & Hooten 2016). Thus, pooling all the available presence data and then combining with a random sample of background pseudo-absences is justified in this case for a data-poor rare species (Biddle *et al*. 2021).

Globally, more than half of all raptor species are threatened, largely due to increasing human land use activities, driving habitat loss and degradation (McClure *et al*. 2018). Quantifying baseline biological parameters, such as range extent and population size is key to establishing a solid foundation from which to build effective conservation action (Watson 2018). With the fundamentals of where a given species occurs and how many individuals exist, conservation planning can be more effectively directed to areas of high conservation priority (IUCN 2001; Rodríguez *et al*. 2007). Our results demonstrate that even with geographically biased occurrence datasets, SDMs can inform globally recognised range metrics and baseline population estimates. In the absence of widespread occurrence data for many rare species, our method is a promising spatial tool with widespread applications for many taxa, in particular for those island endemic species facing high extinction risk.

## Acknowledgements

We thank all staff and volunteers from the Philippine Eagle Foundation who conducted fieldwork over the past four decades, including our local forest guards, nest wardens and Indigenous co-researchers. We are grateful to Cornell University for supplying the restricted eBird location data and thank all individuals and organisations who contributed data to the GRIN information system. LJS thanks The Peregrine Fund for providing a post-doctoral research grant and we thank the M.J. Murdoch Charitable Trust for funding. The PEF would like to thank our local government partners across the Philippines, and the following institutions that funded and supported the field surveys and nest monitoring that led to this paper across the years: Mohammed Bin Zayed Conservation Fund, Local Government of Apayao and Calanasan, Disney Conservation, Whitley Fund for Nature, Microwave Telemetry Inc, KoEko, Forest Foundation Philippines, The Peregrine Fund, Direct Aid Program - AusAID, USAID/Phil-Am Fund, USAID/Protect Wildlife, Insular Life Foundation, GIZ-Coseram, Pacific Paints (Boysen) Philippines, Energy Development Corporation, UNDP Global Environment Fund, Italy Debt Swamp/Department of Finance, US Forest Service, San Roque Power Corporation, Cornell Lab of Ornithology, Raptor Resource Project, and the Department of Environment and Natural Resources through the Biodiversity Management Bureau and its regional and local offices (DENR Regions 2, 4, 8, 9, 10, 11, 12, and 13).

## Data Accessibility Statement

Upon acceptance the data that support the findings of this study will be made openly available on the data repository *figshare*

## Conflict of Interest

The authors have no conflict of interest to declare.

## Supplementary Material

### Materials and Methods

#### Species locations

All Philippine Eagles were trapped using either a modified Bal-Chatri (Miranda & Ibanez 2006) or a large bownet baited with domestic rabbit (*Oryctolagus cuniculus*). Two eagles were instrumented with solar-powered Global Positioning System-Global System for Mobile Communications (GPS-GSM) transmitters while four eagles had battery-powered GPS satellite transmitters fitted harnessed with Teflon-coated nylon ribbon backpacks. All birds were marked with aluminium leg bands – the four females with blue bands on their left leg, and the two males with green bands on their right leg. All GPS transmitter harnessing was conducted with a Gratuitous Permit to trap and tag the birds in the presence of a veterinarian as required by the national government of the Philippines. A total of 80,966 fixes were obtained from four adult females and two adult males from April 2013 to September 2021 (Table S1; Fig. S2). We removed all duplicate fixes and applied a 1-km spatial filter to this raw dataset resulting in 325 spatially filtered GPS fixes. We first fitted models with just the nest and community science occurrences which predicted well but model predictions improved (based on expert validation and model metrics) when we added the GPS fixes, capturing those areas being used by adults outside of the nest sites. Conversely, when we fitted the models with just the GPS fixes this also predicted poorly compared to pooling all the available presence-only occurrence data.

#### Habitat Covariates

We downloaded surface reflectance band imagery from the Moderate Resolution Imaging Spectroradiometer (MODIS) product MCD13A2 (1-km resolution) via passive remote optical sensors onboard the NASA Terra and Aqua satellites, which have 1-2 day overpass frequencies on opposing polar orbits. We used 16-day composite imagery acquired on the 1 June 2020 downloaded using the R package MODIStsp (Busetto & Ranghetti 2016). We downloaded imagery from this date as it corresponds with the start of the wet season in the Philippines and peak vegetation greenness, which has low interannual variability and is generally continuous in forested, mountainous areas throughout the year (Perez & Comiso 2014). We used MODIS imagery because of the high overpass frequencies of the Terra and Aqua satellites, which increases the probability of obtaining cloud free images in tropical regions over the 16-day period. All three bands contain spectral reflectance values estimated by target at surface, calibrated with cloud detection and atmospheric corrections.

We used three surface reflectance bands that represent unclassified raw measures of vegetation structure and composition, better able to capture vegetation patterns in SDMs compared to classified remote sensing vegetation products (Morán-Ordóñez *et al*. 2012; Shirley *et al*. 2013), including for tropical forest taxa (Van doninck *et al*. 2020). Band 1 Red represents photosynthetic activity related to plant biomass, Band 2 Near Infrared represents leaf and canopy structure, with B7 Short Wave Infrared related to senescent or old growth biomass (Shirley *et al*. 2013). Reflectance values are expressed as the ratio of reflected over incoming radiation, meaning reflectance can be measured between the values of zero and one. Absolute reflectance values of 3-4 indicate healthy vegetation (Huete *et al*. 2004).

Evergreen Forest is a consensus product of percentage land cover integrating GlobCover (v2.2), MODIS land-cover product (v051), GLC2000 (v1.1) and DISCover (v2) (Tuanmu & Jetz 2014), used here to represent dipterocarp forest. Human Footprint Index (HFI) represents human population density, land use and infrastructure, including built-up areas and access routes such as roads and rivers (WCS, CIESIN 2005), used here as we expect Philippine Eagles to avoid areas of high human impact. Leaf Area Index (LAI) is a biophysical measure of the amount of foliage within the plant canopy based on MODIS vegetation products, used here as a composite Dynamic Habitat Index (DHI) product spanning the years 2003-2014 (Radeloff 2019). LAI values range from 0 (bare ground) to > 10 (dense coniferous forest) and is a key driver of primary productivity (Asner *et al*. 2003). The DHI product summarises three measures of vegetation productivity: annual cumulative, minimum throughout the year, and seasonality as the annual coefficient of variation. Combined, we used the LAI Dynamic Habitat Index as a proxy for food availability, assuming that higher LAI values would be associated with higher species richness (Hobi *et al*. 2017). All selected covariates showed low collinearity (Variance Inflation Factor (VIF) < 4; Table S3).

#### Species Distribution Models

In its original implementation MAXENT imposed a ‘lasso’ (least absolute shrinkage and selection operator) regularization penalty, where only the most significant covariates are retained, with uninformative covariates set at zero. Instead, the maxnet package uses an elastic net penalty (via the glmnet package, Friedman *et al*. 2010) to perform automatic covariate selection (lasso) and continuous shrinkage (ridge regression) simultaneously (Zou & Hastie 2005; Phillips *et al*. 2017), evaluating the contribution of all covariates and shrinking low-contribution coefficients towards zero. Elastic net regularization improves predictive accuracy compared to the lasso, in both simulated and real data examples (Zou & Hastie 2005) and may be viewed as a generalization of the lasso. Overall, penalizing model coefficients reduces model variance, resulting in a regression model that generalizes better (Valavi *et al*. 2021). We parametrized the penalized logistic regression model using infinite weighting (presence weights = 1, background = 100), within the inhomogeneous Poisson process framework because this is the most effective method to model presence-background data as used here (Warton & Shepherd 2010; Hefley & Hooten 2015).

Within the maxnet package the complementary log-log (cloglog) link function was selected as a continuous index of habitat suitability, with 0 = low suitability and 1 = high suitability. Phillips *et al*. (2017) demonstrated the cloglog link is equivalent to an inhomogeneous Poisson process and can be interpreted as a measure of relative occurrence probability proportional to a species potential abundance. Optimal-model selection was based on Akaike’s Information Criterion (Akaike 1974) corrected for small sample sizes (AIC_c_; Hurvich & Tsai 1989), to determine the most parsimonious model from two maxnet parameters: regularization beta multiplier (β; level of coefficient penalty) and feature classes (response functions, Warren & Seifert 2011; Phillips *et al*. 2017). Thirty-eight candidate models of varying complexity were built by conducting a grid search with a range of regularization multipliers from 1 to 10 in 0.5 increments, and two feature classes (response functions: Linear, Quadratic) in all possible combinations using the ‘block’ method of spatial cross-validation (*k =* 5) in the ENMeval package in R (Muscarella *et al*. 2014).

Spatial block cross-validation divides the geographical structure of the data according to latitudinal and longitudinal lines, dividing all occurrences into four spatially independent bins of equal numbers. By binning the geographical structure of test data into blocks, the models are projected onto an evaluation region not included in the calibration process. All occurrence and background test points are assigned to their respective bins dependent on location, thus further reducing spatial auto-correlation between testing and training localities (Muscarella *et al*. 2014). We considered all models with a ΔAIC_c_ < 2 as having strong support (Burnham & Anderson 2004) and selected the model with the lowest ΔAIC_c_ that used both feature classes (Linear, Quadratic) with the highest regularization penalty and omission rate closest to 0.10 (OR10).

#### Population size estimation

For calculating the median habitat area and population size estimates we fitted Kernel Density Estimators (KDE) using a 95 % bivariate normal kernel with an ad- hoc reference smoothing parameter (*h_ref_* ). We then fitted 99 % Minimum Convex Polygons (MCP) to calculate the maximum habitat area, and thus minimum population size, and Local Convex Hulls (LoCoH) using the sphere of influence radius method (r-LoCoH, maximizing all nearest neighbour distances (Getz *et al*. 2007)), for calculating the minimum habitat area and thus maximum population size. We summed all individual home range estimates from each adult and calculated a median value for each estimator method. We then used the three median values from each home range estimator to apply an overall median, and minimum to maximum range for calculating population sizes. Because of varied telemetry sampling rates between the six adults, we subsampled the raw location fixes using a minimum 3-hour interval between fixes to achieve consistency across individual estimators. All home range estimates were calculated in the R package adehabitatHR (Calenge 2006), using code adapted from Tétreault & Franke (2017).

## Supplementary Tables

**Table S1.** GPS metadata for the six tagged adult Philippine Eagles from the island of Mindanao, used for home range estimation. Fixes are subsampled from the raw data locations using a 3-hr sampling rate interval.

| ID | From | To | Fixes |
| --- | --- | --- | --- |
| 001F | 16/02/2014 | 10/05/2015 | 1,186 |
| 002F | 22/12/2014 | 20/01/2016 | 1,063 |
| 003F | 11/04/2013 | 19/02/2014 | 263 |
| 004M | 19/04/2014 | 05/08/2014 | 190 |
| 005M | 17/11/2019 | 12/09/2021 | 5,252 |
| 006F | 15/10/2019 | 05/06/2021 | 2,344 |
| Total |  |  | 10,298 |

**Table S2.** Model selection metrics for all six candidate models with ΔAIC_c_ < 2. RM = regularization multiplier (β), FC = feature classes, LQ = Linear, Quadratic, OR10 = 0.10 omission rate.

| Model | RM | FC | AIC<sub>c</sub> | $\Delta AIC_c$ | OR10 |
| --- | --- | --- | --- | --- | --- |
| 1 | 1.0 | L | 2083.954 | 1.832 | 0.117 |
| 2 | 1.0 | LQ | 2082.122 | 0.000 | 0.128 |
| 3 | 1.5 | L | 2083.970 | 1.848 | 0.117 |
| 4 | 1.5 | LQ | 2083.786 | 1.665 | 0.117 |
| 5 | 2.0 | L | 2084.006 | 1.885 | 0.117 |
| 6 | 2.5 | L | 2084.072 | 1.950 | 0.117 |

**Table S3.** Multi-collinearity test using stepwise elimination Variance Inflation Factor (VIF) analysis. Covariates with VIF < 4 have low correlation with other covariates, and thus are suitable for inclusion in calibration models when further evaluated for ecological relevance.

| Covariate | VIF |
| --- | --- |
| Band 1 Red | 3.14 |
| Band 7 Shortwave infrared | 3.10 |
| Band 2 Near infrared | 1.83 |
| Evergreen Forest | 1.67 |
| Leaf Area Index | 1.30 |
| Human Footprint Index | 1.27 |

## Supplementary Figures

**Figure S1.**
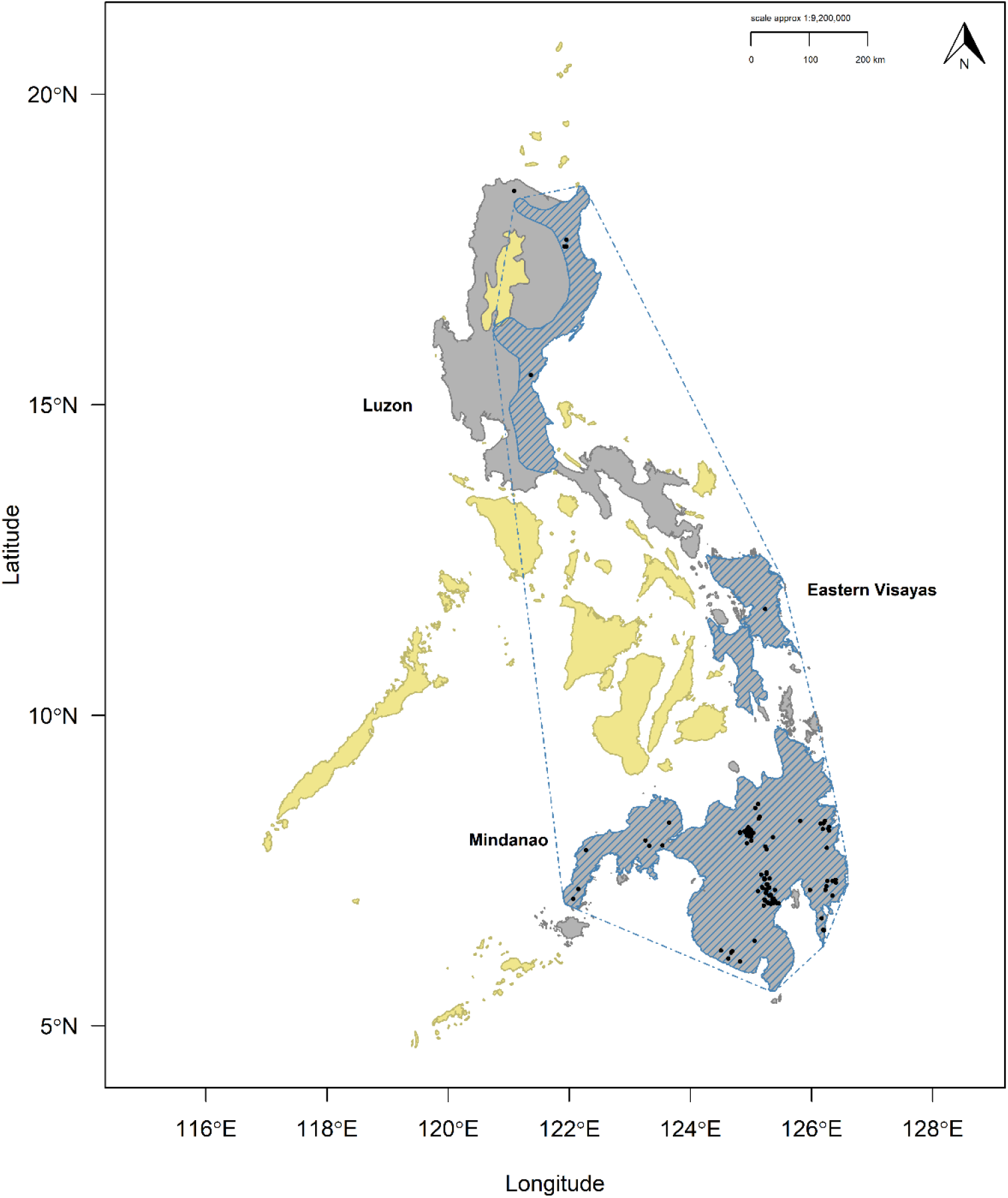
Range map for the Philippine Eagle with our model accessible area (dark grey) and IUCN range map (hashed blue areas) and IUCN EOO polygon (hashed blue line). Yellow polygons define the national boundary of the Philippines outside of the species accessible area. Black points define unfiltered Philippine Eagle occurrences from nests and community science data. For clarity, GPS locations from the six tagged adults in Mindanao are shown in Fig. S2.

**Figure S2.**
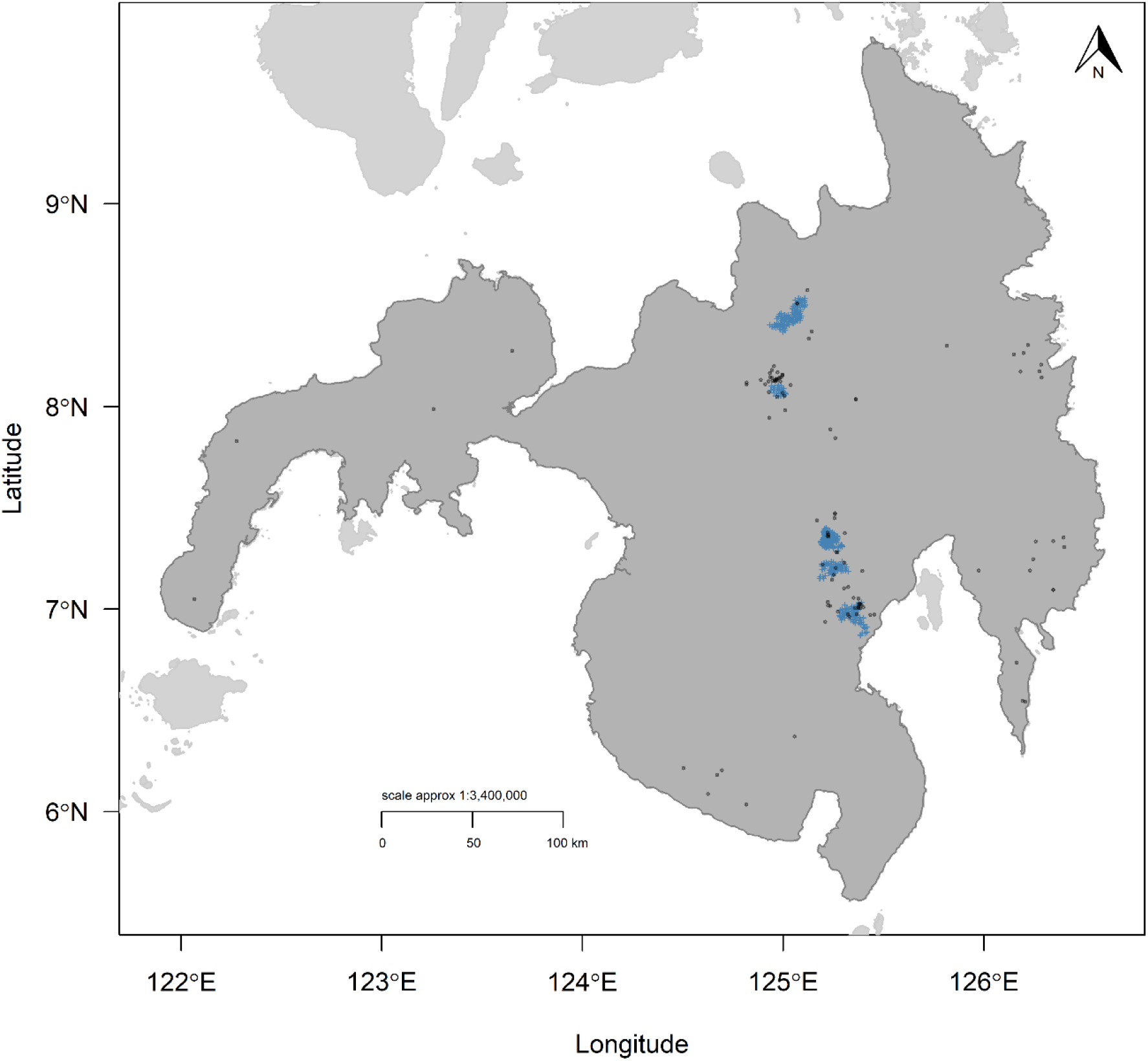
Filtered GPS fixes using a 1-km filter (blue points) for the six tagged adult Philippine Eagles from the island of Mindanao, used in the Species Distribution Models. Black points denote the Philippine Eagle occurrences from nests and community science data.

**Figure S3.**
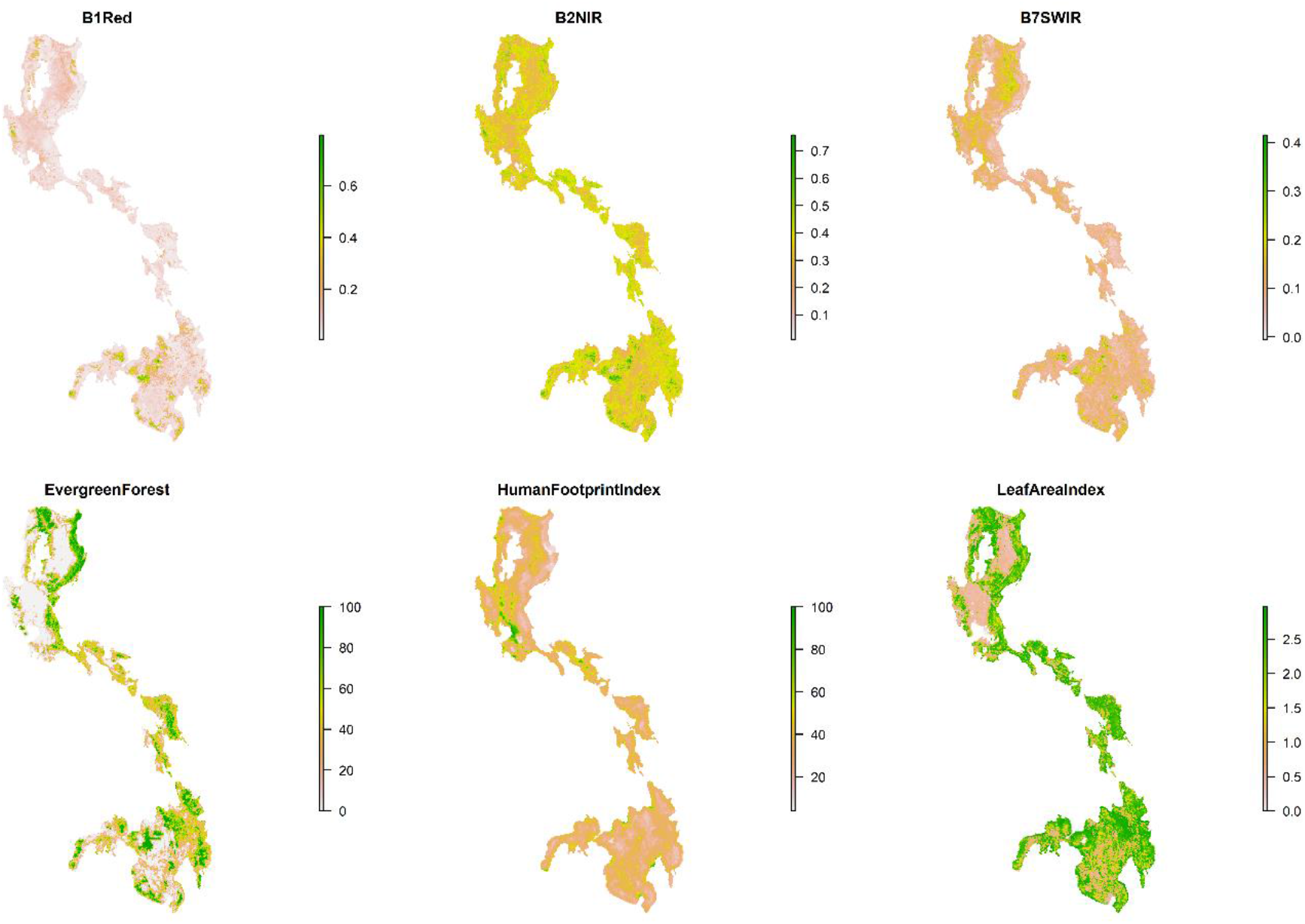
Habitat covariates used in Species Distribution Models for the Philippine Eagle.

**Figure S4.**
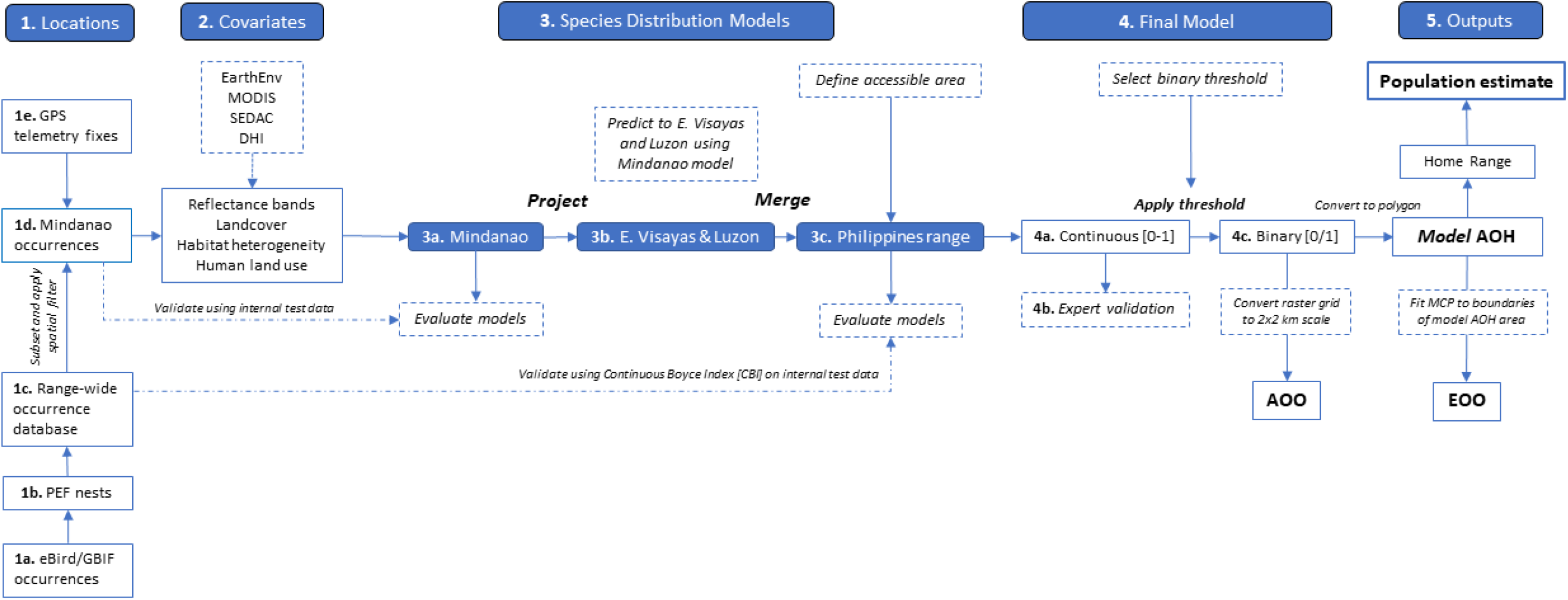
Spatial model workflow for the Philippine Eagle Species Distribution Model to estimate range metrics and population size.

**Figure S5.**
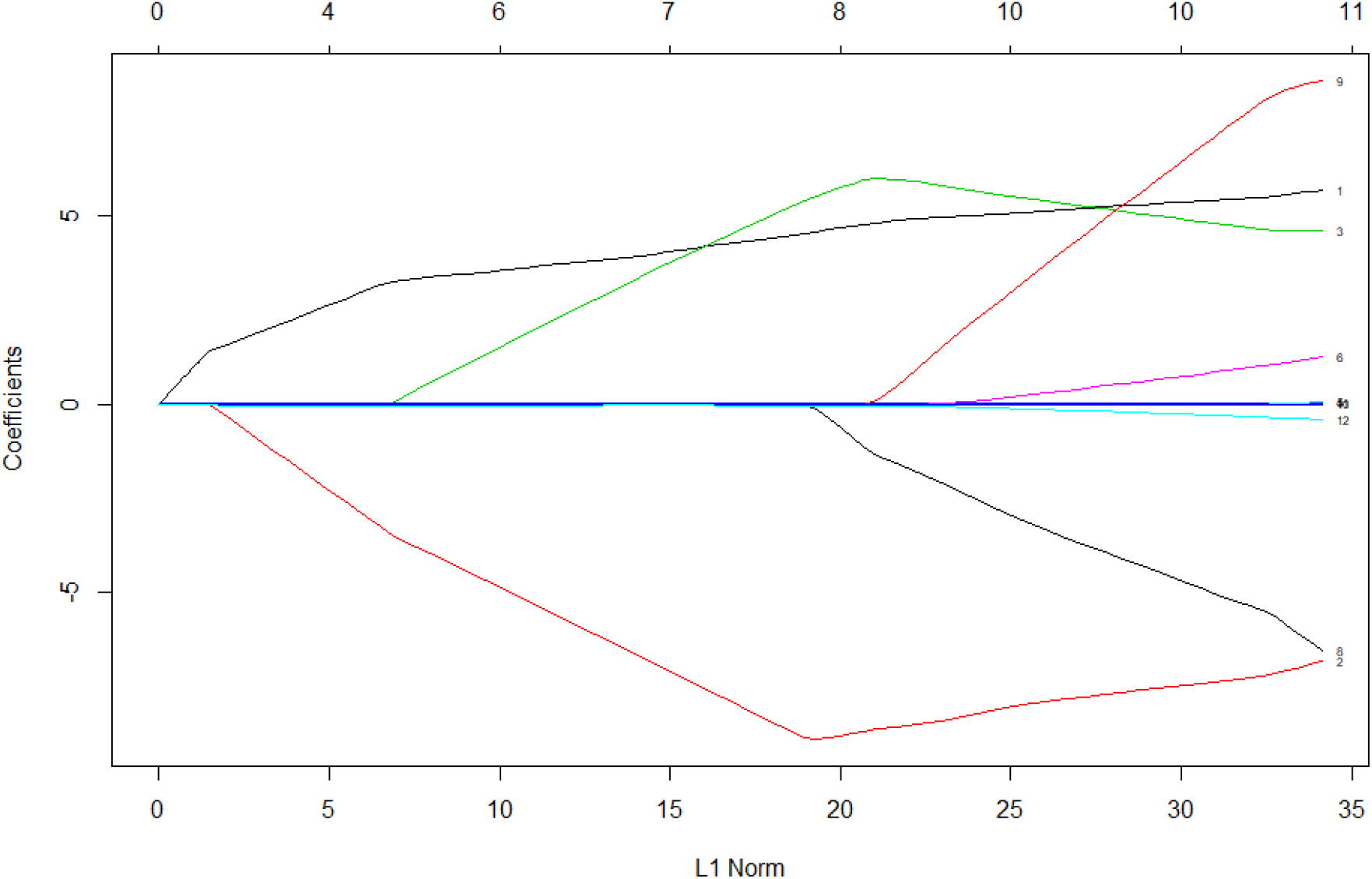
Beta coefficient paths for the optimal penalized logistic regression model where each curve corresponds to a covariate term (linear and quadratic). The paths of each coefficient term are plotted against the L1-norm (lasso or elastic net) of the whole coefficient vector as lambda (the amount defining the level of coefficient shrinkage) varies. The upper axis indicates the number of non- zero coefficients at the current lambda which is the effective degrees of freedom for the lasso or elastic net.

**Figure S6.**
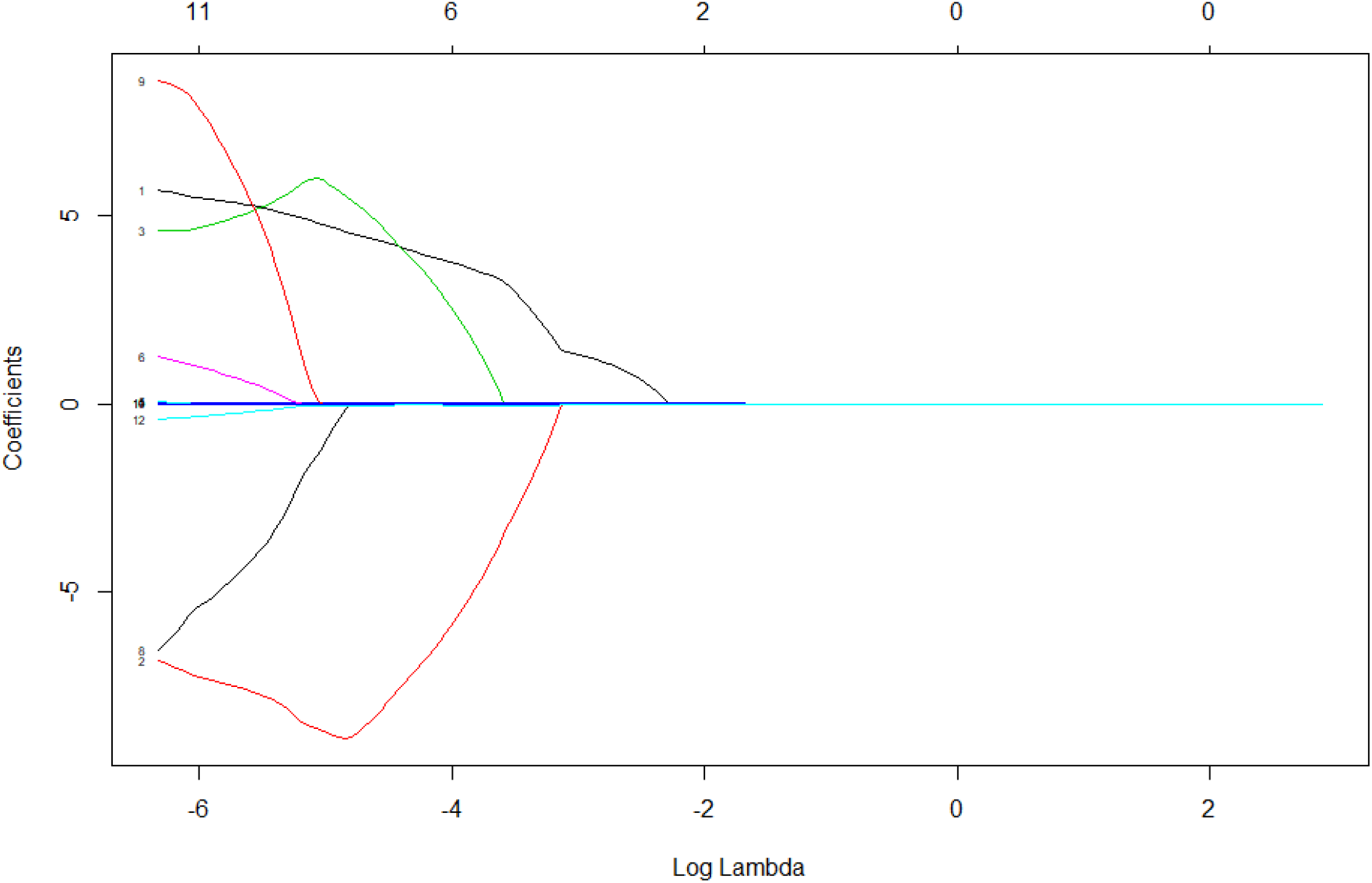
Beta coefficient paths for the optimal penalized logistic regression model where each curve corresponds to a covariate term (linear and quadratic). The paths of each coefficient term are plotted against the log-lambda of the whole coefficient vector as lambda (the amount defining the level of coefficient shrinkage) varies. Log-lambda on the y-axis indicates the log of the optimal value of lambda which minimizes the prediction error. This lambda value will give the most accurate model. The upper axis indicates the number of decreasing non-zero coefficients at the current lambda which is the effective degrees of freedom for the elastic net.

**Figure S7.**
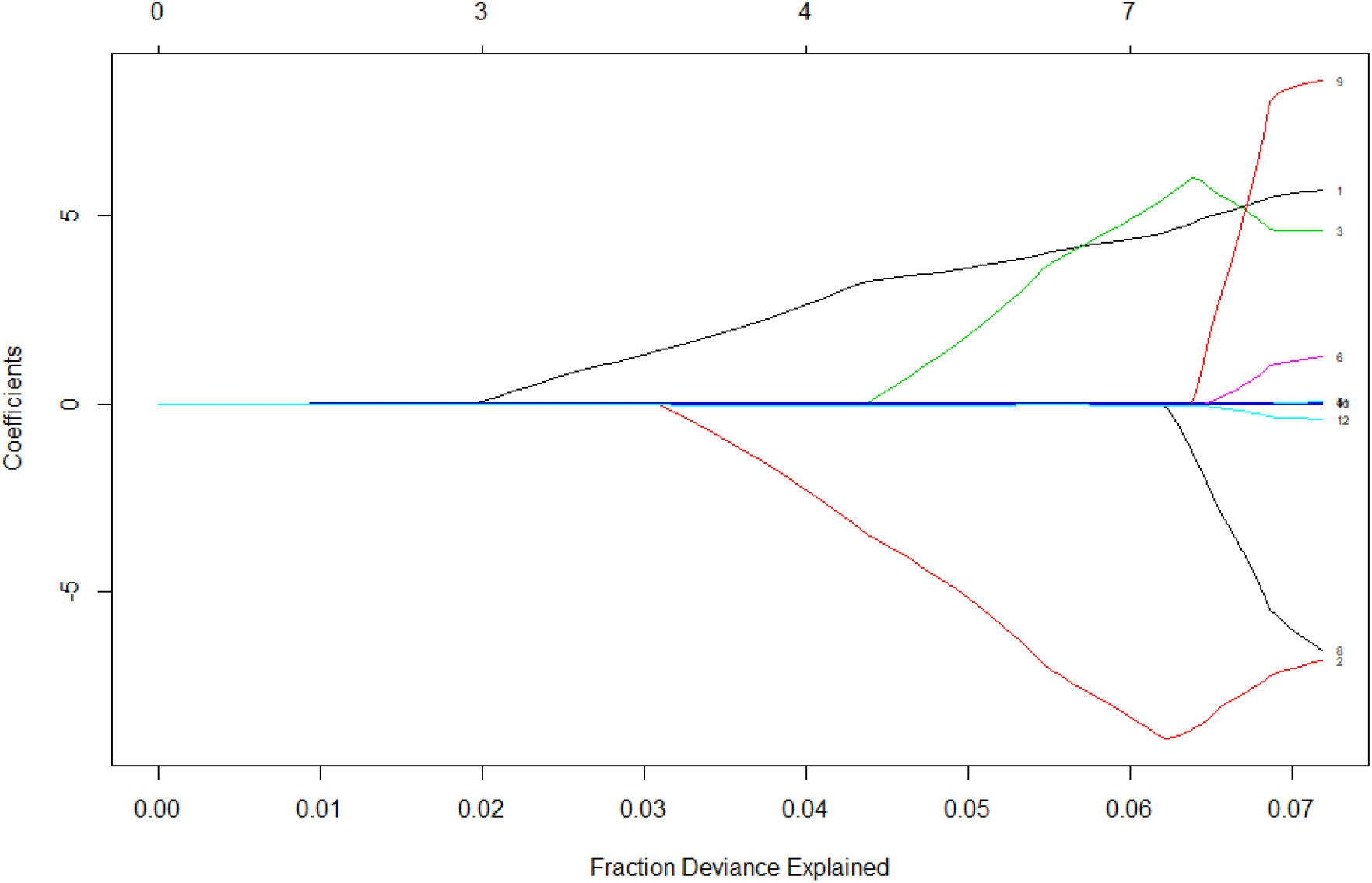
Beta coefficient paths for the optimal penalized logistic regression model where each curve corresponds to a covariate term (linear and quadratic). The paths of each coefficient term are plotted against the fraction deviance explained on the training data. The upper axis indicates the number of non-zero coefficients at the current lambda which is the effective degrees of freedom for the elastic net.

**Figure S8.**
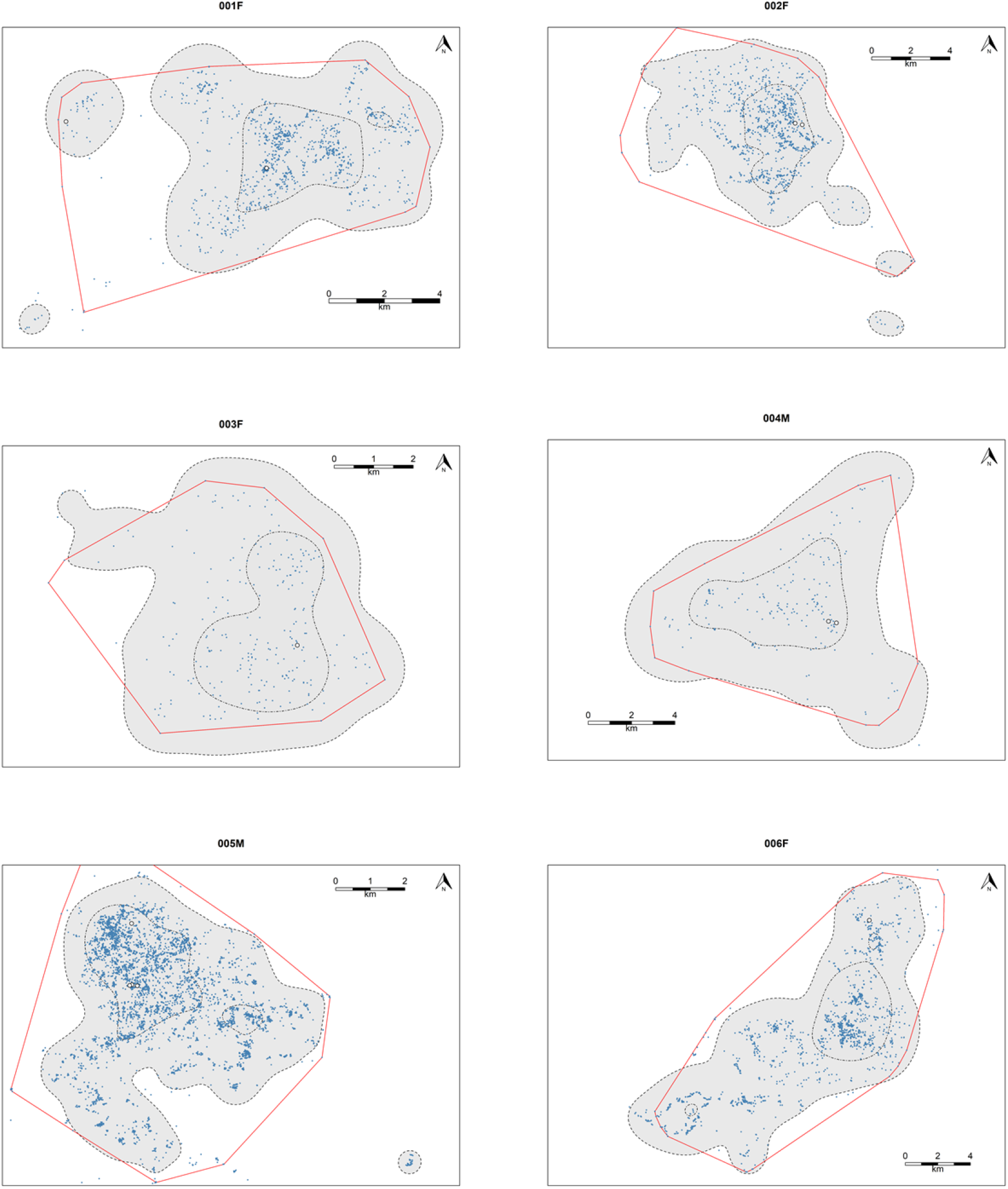
Home range estimates for six adult Philippine Eagles using a bivariate 95% Kernel Density Estimate (KDE, light grey with black dashed line) and 99% Minimum Convex Polygon (MCP, red line). Blue points are GPS fixes, white points known nests, with 50% KDE estimate shown in black dot-dash line.

**Figure S9.**
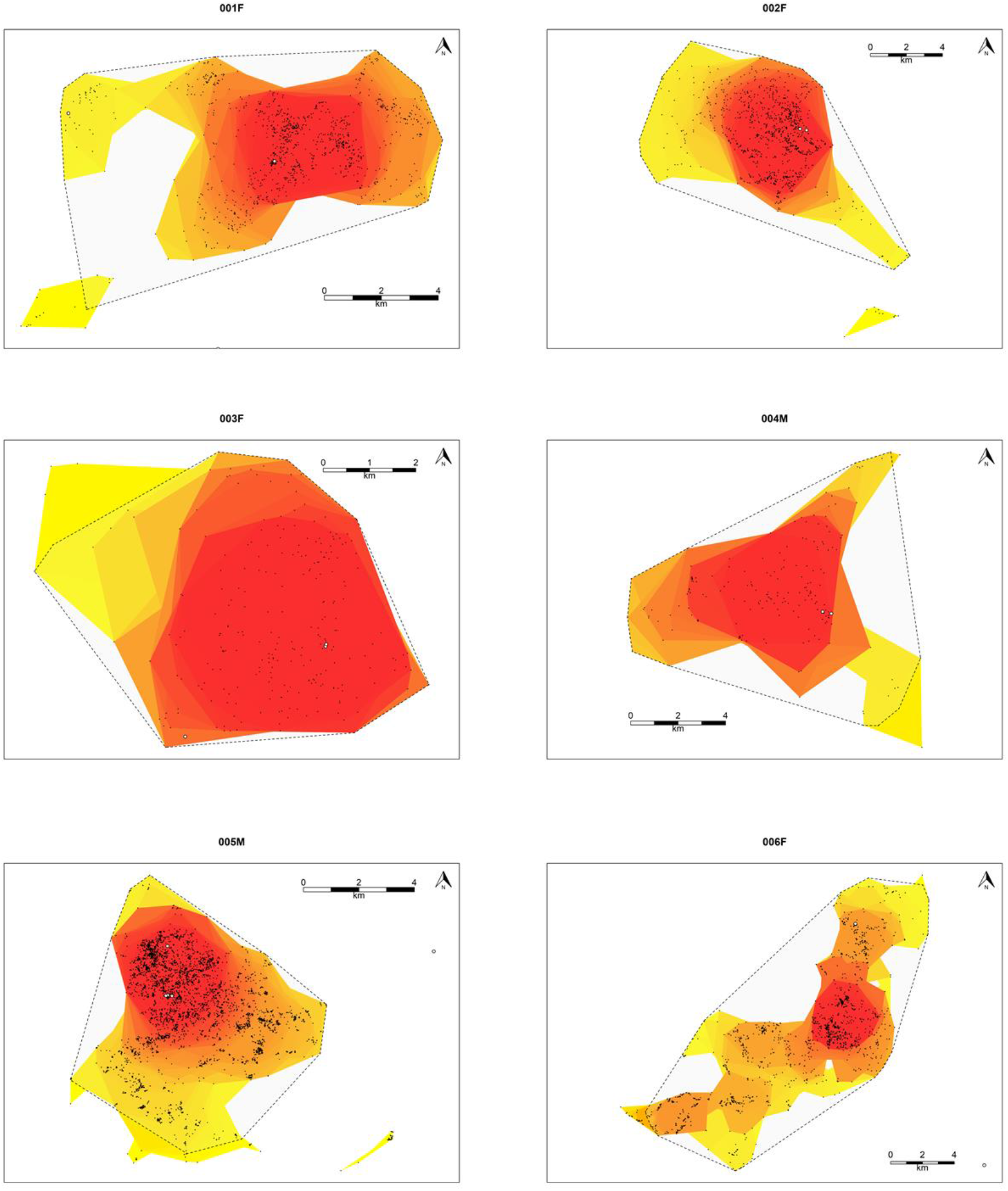
Home range estimates for six adult female Philippine Eagles using a radius Local Convex Hull estimator (r-LoCoH) and 99% Minimum Convex Polygon (MCP, black hashed line). Black points are GPS fixes, white points known nests. The gradient for utilization distribution is represented from high use (red) to low use (yellow).

**Figure S10.**
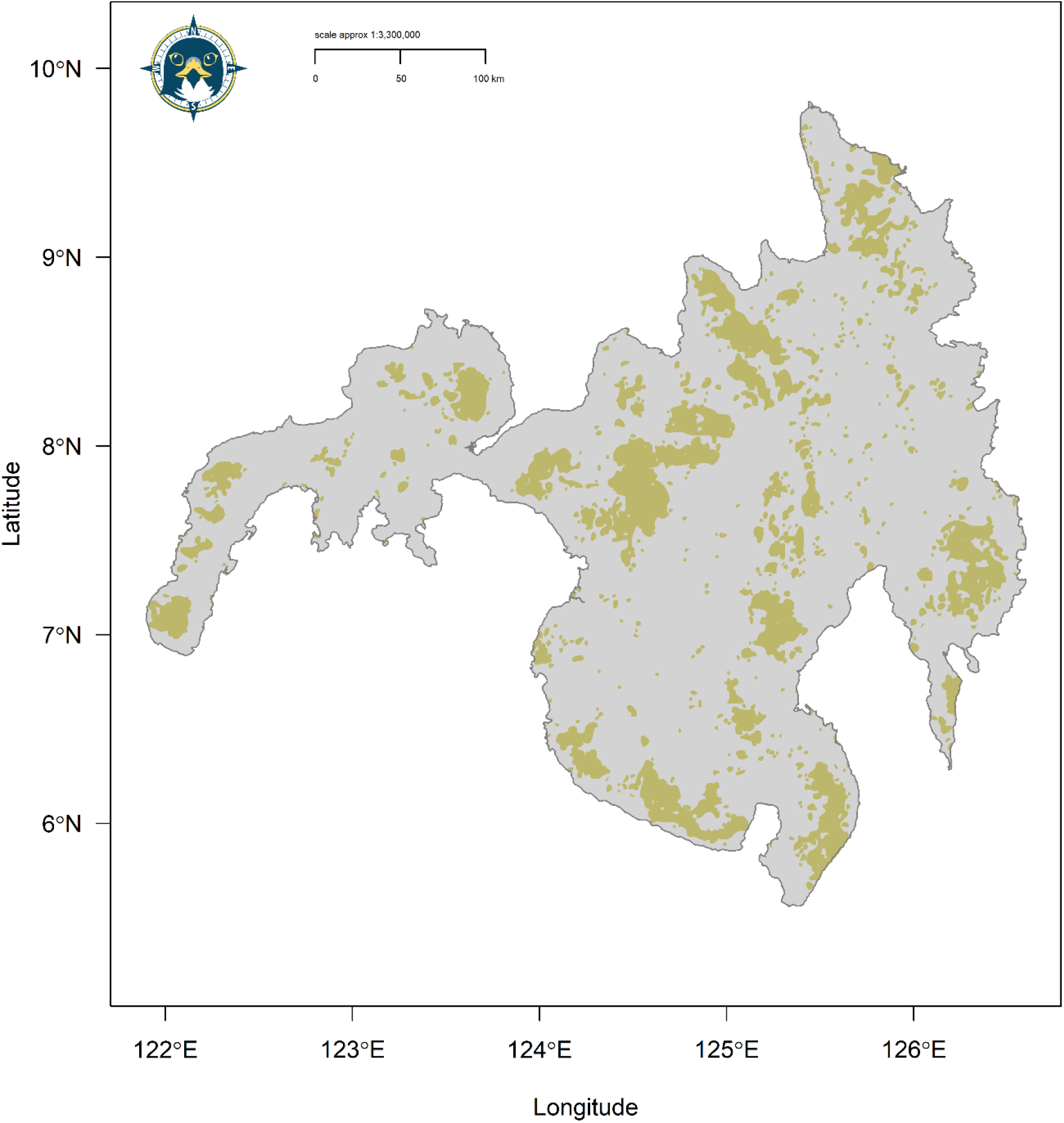
Reclassified binary *model* AOH area (dark khaki) for the Philippine Eagle on the island of Mindanao.

**Figure S11.**
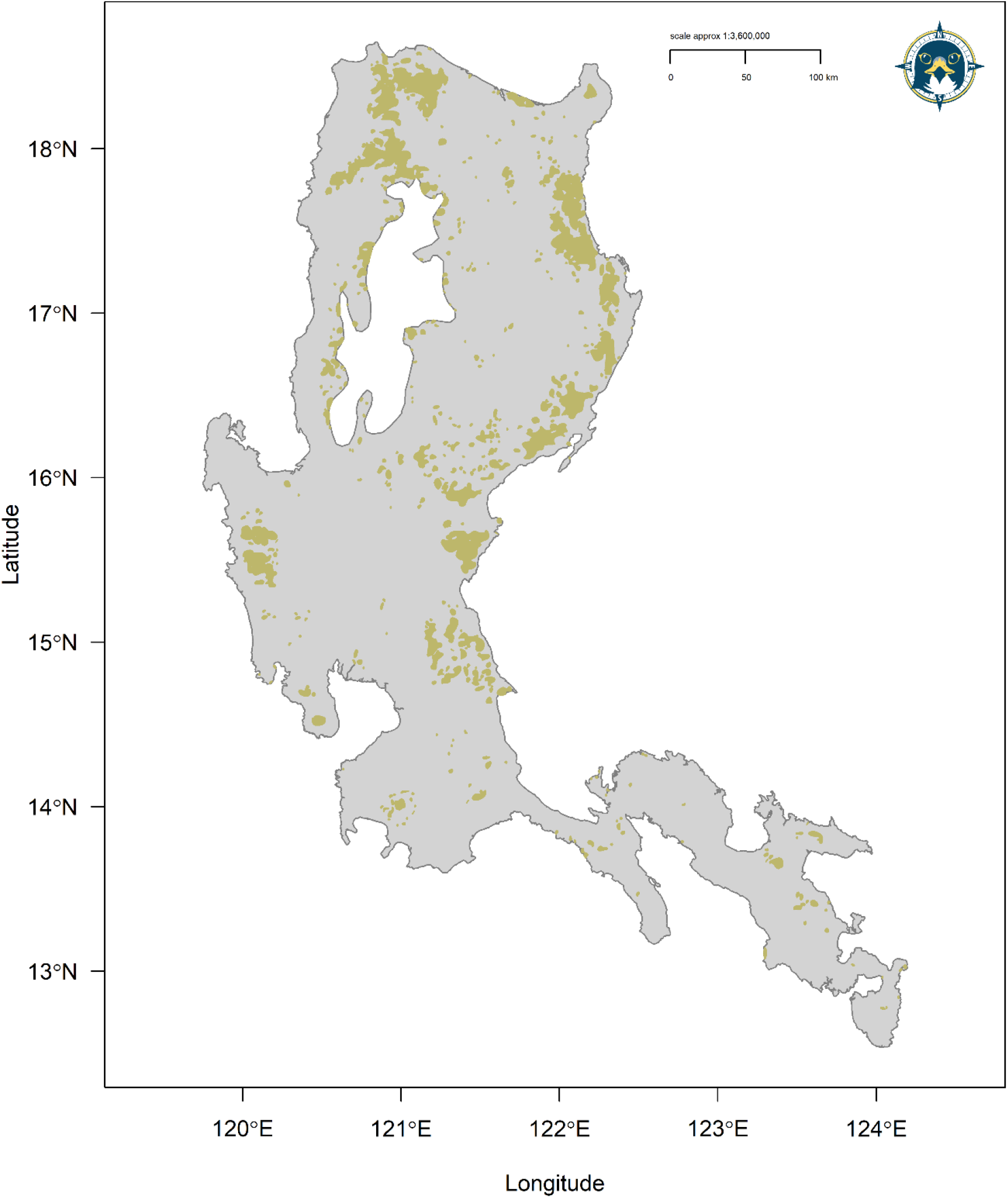
Reclassified binary *model* AOH area (dark khaki) for the Philippine Eagle on the island of Luzon.

**Figure S12.**
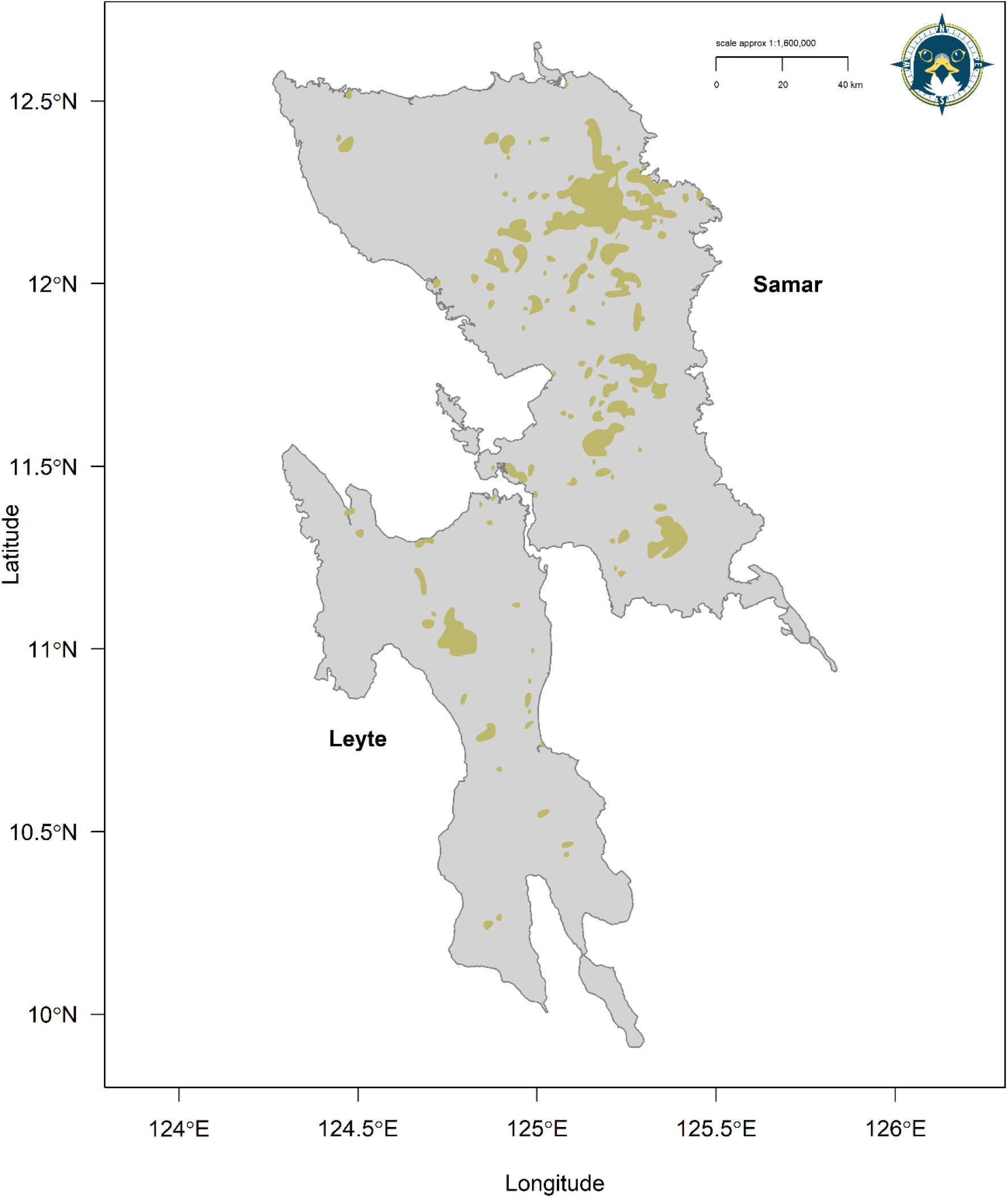
Reclassified binary *model* AOH area (dark khaki) for the Philippine Eagle on the islands of Leyte and Samar in the Eastern Visayas.

**Figure S13.**
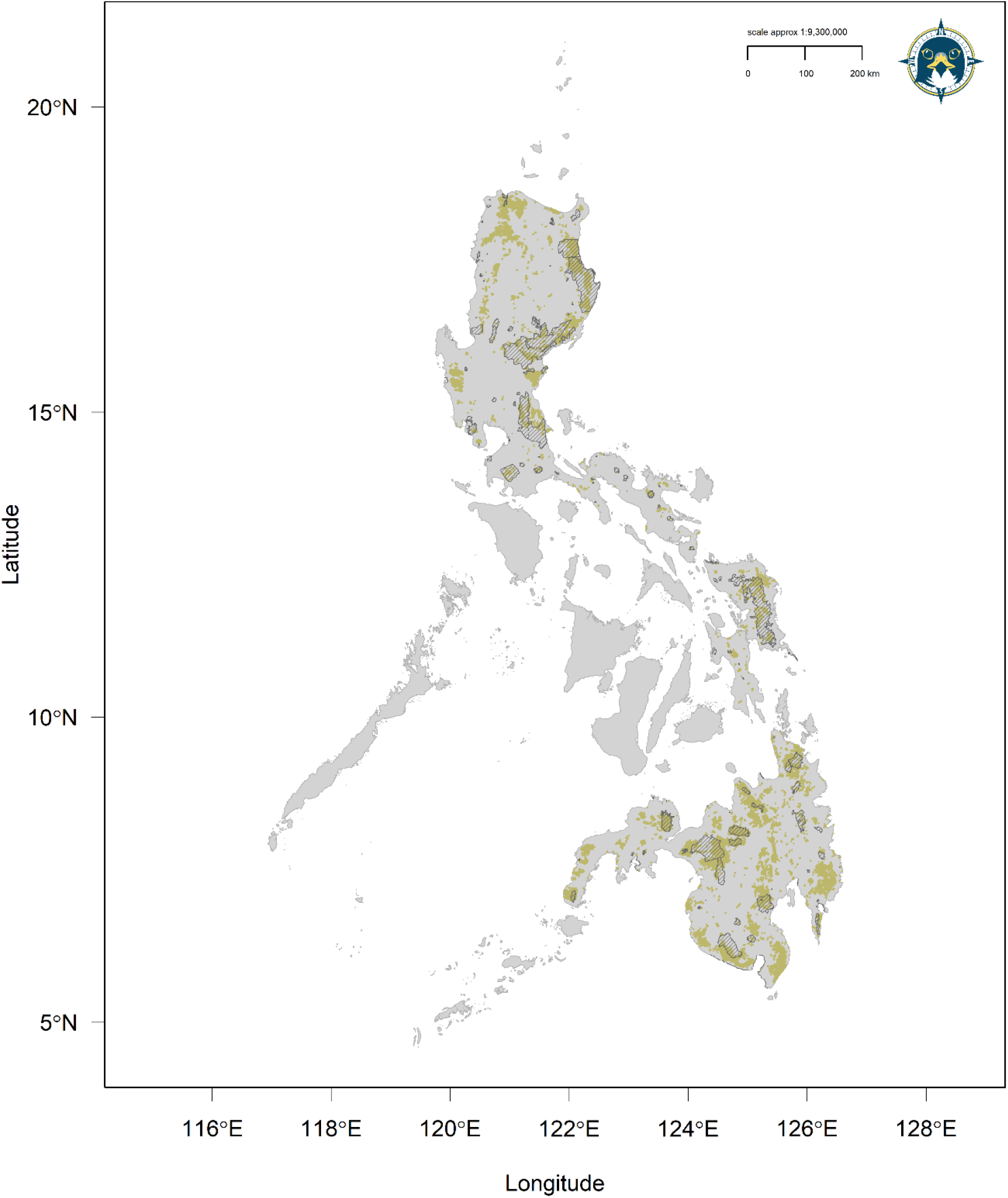
Reclassified binary *model* AOH area (dark khaki) for the Philippine Eagle showing spatial coverage of the World Database on Protected Areas (WDPA) network (grey polygons) compared to the *model* AOH polygon area.

**Figure S14.**
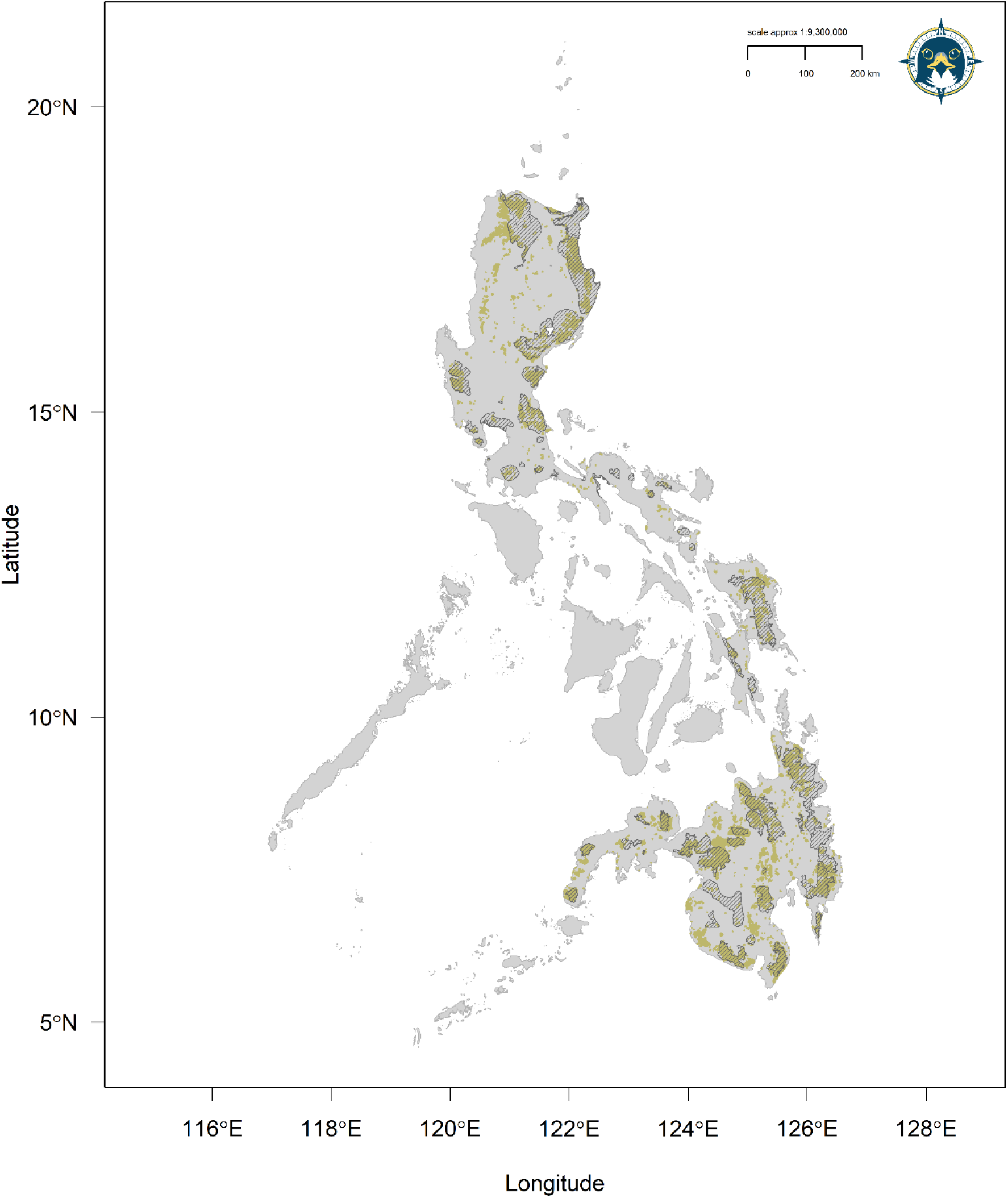
Reclassified binary *model* AOH area (dark khaki) for the Philippine Eagle showing spatial coverage of the Key Biodiversity Area (KBA) network (grey polygons) compared to the *model* AOH polygon area.

## Notes

### Competing Interest Statement

The authors have declared no competing interest.

### Summary of Updates

New species distribution models fitted using remote sensing covariates from surface reflectance values, which were then used to recalculate range metrics, population size and a protected area spatial gap analysis.

